# Characterization of Electrophysiological and Transcriptomic Alterations in Patient-Derived Neurons from CHAMP1 Syndrome

**DOI:** 10.64898/2026.06.19.733368

**Authors:** Dailey Nettles, Christina Stanton, Zach Hunter, Bryan Granger, Elizabeth Wallace, Andrew Lutsky, Suganya Subramanian, Meris Privette, Lori McMahon, Stefano Berto

## Abstract

Mutations in chromosome alignment maintaining phosphoprotein 1 (CHAMP1) have been linked to neurodevelopmental disorders characterized by intellectual disability, developmental delay, and autism spectrum disorder. However, the cellular and electrophysiological mechanisms by which CHAMP1 mutations disrupt human neuronal development remain poorly understood. In the present study, we used patient-derived induced pluripotent stem cells (iPSCs) carrying two pathogenic CHAMP1 mutations and generated neural progenitor cells (NPCs) and excitatory neurons to investigate the effects of each mutation on neuronal maturation and function, DNA repair, and gene expression. Proliferative capacity declines with CHAMP1 dosage, while DNA repair dysfunction is allele-specific. Whole-cell patch-clamp electrophysiology revealed that CHAMP1 mutant neurons exhibit significant alterations in intrinsic membrane properties during early developmental stages, including depolarized resting membrane potential, reduced action potential firing, and impaired waveform kinetics. These functional deficits were accompanied by reduced sodium and potassium current densities, suggesting impaired ion channel accumulation during neuronal maturation. Furthermore, recordings of spontaneous excitatory postsynaptic currents indicated altered synaptic activity and reduced proportions of synaptically active neurons. Morphological analyses showed that CHAMP1-deficient neurons exhibit impaired neurite outgrowth and branching, supporting a defect in neuronal maturation. Single-nucleus transcriptomic profiling further revealed delayed developmental trajectories and mutation-specific dysregulation of synaptic gene programs enriched for autism, ADHD, and epilepsy risk genes. Together, these findings demonstrate that CHAMP1 mutations disrupt multiple aspects of neuronal development, including homologous recombination (HR) dysfunction in NPCs, membrane excitability, ion channel function, and synaptic connectivity. Our results provide insights into the neurobiological consequences of CHAMP1 mutations and establish patient-derived neurons as a platform to investigate cellular pathophysiology and potential therapeutic strategies for CHAMP1-associated neurodevelopmental disorders.

## INTRODUCTION

*Chromosome alignment maintaining phosphoprotein 1 (CHAMP1)* encodes a zinc finger protein that regulates chromosome segregation and kinetochore–microtubule attachments during mitosis^1,2^. Recent prospective phenotyping and large-scale sequencing studies have expanded the clinical spectrum associated with CHAMP1 variants, encompassing global developmental delay (GDD), intellectual disability (ID), microcephaly, hypotonia, and dysmorphic features, together with prominent neurobehavioral involvement: autism spectrum disorder (ASD) in approximately one-third and attention-deficit/hyperactivity disorder (ADHD) in roughly 60% of individuals, accompanied by repetitive behaviors, sensory-seeking symptoms, and, in some cohorts, epilepsy, sleep disturbance, and gastrointestinal abnormalities^3–6^. Pathogenic CHAMP1 variants are predominantly de novo and protein-truncating but also include whole-gene deletions, as part of 13q34 deletion syndrome, and rarer missense changes. Recent work directly comparing these variant classes found that individuals carrying truncating mutations exhibit significantly more severe impairment across adaptive, developmental, and medical domains than those with whole-gene deletions, suggesting that truncating mutations act through dominant-negative or gain-of-function mechanisms, whereas deletions act through CHAMP1 haploinsufficiency^7^. Given that different classes of CHAMP1 variants may perturb protein function through distinct, and potentially opposing, mechanisms, it remains unclear whether they converge on similar, distinct, or overlapping cellular and clinical phenotypes.

Mechanistically, CHAMP1 localizes to the kinetochore, the structure that mediates attachment between centromeres and spindle microtubules and supports the spindle assembly checkpoint during metaphase^8–10^. This function is particularly relevant during early cortical development, when human apical progenitors exhibit a prolonged metaphase relative to other species^11,12^. Disruption of accurate chromosome segregation can impair neuronal proliferation, increase chromosomal abnormalities, and promote apoptosis, thereby altering brain development^13–15^. Consistent with this, patient-derived lymphoblasts carrying CHAMP1 mutations show abnormal spindle formation, and CHAMP1 depletion in HeLa cells disrupts cytokinesis, proliferation, and migration^16^. Beyond its mitotic role, CHAMP1 interacts with POGZ and MAD2L2 (REV7) through its C-terminus to influence homologous recombination repair and DNA damage response pathway choice^17–19^. Because pathogenic truncating variants are predicted to disrupt this C-terminal region, they may compromise these interactions in a manner distinct from gene deletion. In vivo, CHAMP1 loss alters brain size and behavior, including learning, memory, social interaction, and depression-like behavior, and CHAMP1 knockdown delays neuronal differentiation in mouse cultures, with concomitant dysregulation of POGZ expression indicating functional coupling between the two genes^19^. Despite this strong genetic and mechanistic link between CHAMP1 and neurodevelopmental disorders, how CHAMP1 dysfunction disrupts human neural development remains poorly understood.

Human induced pluripotent stem cell (hiPSC)-derived neurons from individuals with CHAMP1 syndrome provide a powerful system to interrogate disease-relevant neuronal mechanisms. These patient-specific models recapitulate key features of early human neurodevelopment in a controlled in vitro context, enabling detailed investigation of mutation-specific effects on neuronal differentiation, morphology, synaptic function, and electrophysiological properties^20–26^. More broadly, hiPSC-derived neuronal systems have become an essential platform for modeling rare genetic and neurodevelopmental disorders (NDD), offering otherwise inaccessible access to human neuronal tissue^27–29^.

In this study, we leveraged patient-derived and isogenic hiPSC models to define CHAMP1 function across neural progenitor and neuronal stages. We integrated assays of DNA damage repair, cellular proliferation, electrophysiology, neuronal morphology, and single-nucleus transcriptomics. Our results demonstrate that CHAMP1 mutations compromise homologous recombination and reduce proliferative capacity in neural progenitors, while also disrupting the timing and maturation of excitatory neurons. At the functional level, these defects manifest as altered membrane properties, reduced action potential firing, decreased sodium current density, impaired synaptic activity, and aberrant neurite outgrowth. Single-nucleus transcriptomic profiling links these phenotypes to a coordinated molecular program as CHAMP1-mutant cells follow delayed developmental trajectories and show mutation-specific yet convergent dysregulation of gene networks.

To our knowledge, this is the first study to systematically examine CHAMP1 variants in models of the developing human brain, integrating electrophysiological and transcriptomic signatures that may underlie patient phenotypes and represent candidate therapeutic biomarkers. Collectively, our findings establish CHAMP1 as a critical, multi-level regulator of human neurodevelopment, linking genome stability in progenitors to the transcriptional timing and synaptic maturation of neurons and providing mechanistic insight into how different pathogenic variants drive neurodevelopmental disease.

## METHODS

### Study design

This study investigated the cellular, electrophysiological, and transcriptomic consequences of CHAMP1 mutations in patient-derived neural progenitor cells (NPCs) and neurons, focusing on early developmental stages, proliferation, cell-cycle regulation, and differentiation prior to synaptic maturation. Given CHAMP1’s roles in kinetochore function and chromosome segregation, we prioritized assays of neuronal maturation and function, neurite arborization, dendritic morphology, and single-nucleus transcriptomics. To ensure rigor and reproducibility, all experiments used at least three independent biological replicates derived from distinct hiPSC differentiations, and cellular analyses quantified ≥20 NPCs or neurons per replicate to provide adequate statistical power.

### Human iPSCs

Four hiPSC lines were used in this study (Table S1): one control line, two patient-derived lines, and one isogenic knockout. The control line (K3), referred to here as CHAMP1⁺ᐟ⁺, was derived from a neurotypical 1-month-old male and obtained from the Duncan laboratory (Department of Regenerative Medicine and Cell Biology, Medical University of South Carolina). The control line (20b), referred to here as CHAMP1_2⁺ᐟ⁺, was derived from a neurotypical 55 years-old male^30^ and obtained from the Lizarraga laboratory (Brown University). Two patient-derived lines were obtained from the Coriell Institute for Medical Research: CHAMP1^Arg497*^ (Coriell #GM27860), carrying a nonsense mutation, and CHAMP1^Ile698Metfs*5^ (Coriell #GM29104), carrying a frameshift mutation; these are referred to as CHAMP1A and CHAMP1B, respectively. An isogenic CHAMP1 knockout line (iCHAMP1⁻ᐟ⁻) was generated on the K3 (CHAMP1⁺ᐟ⁺) background (see below). All iPSC colonies were inspected daily to confirm the absence of spontaneous differentiation and were maintained in mTeSR Plus basal medium supplemented with 5× supplement (STEMCELL Technologies, #100-0276) on Geltrex-coated plates (Gibco, #A1413202). Cells were fed daily and passaged every 4–5 days at a 1:6 ratio using ReLeSR (STEMCELL Technologies, #100-0483). To minimize the accumulation of chromosomal abnormalities, iPSCs were maintained only up to passage 50. Pluripotency was periodically confirmed by immunostaining for core markers, including OCT4 (Abcam, #Ab19857), TRA-1-60 (Abcam, #Ab16288), and NANOG (Abcam, #Ab109250).

### Generation of the isogenic CHAMP1 knockout (iCHAMP1⁻*ᐟ*⁻) line

Isogenic CHAMP1 knockout (KO) human induced pluripotent stem cell (iPSC) lines were generated using CRISPR-Cas9 genome editing. Exon 3 of CHAMP1 was targeted using one of two single-guide RNAs (sgRNAs) designed with Benchling: sgRNA1, 5′-CAATGTTCAGGAGACAACGT-3′ (PAM: GGG), or sgRNA2, 5′-TCTCCAGAGCTTCGCAAACC-3′ (PAM: CGG). sgRNA sequences were cloned into the PX459 v2.0 plasmid encoding SpCas9 and puromycin resistance (Addgene plasmid #62988). Plasmids were transfected into iPSCs using ViaFect Transfection Reagent (Promega, E4981). Twenty-four hours after transfection, cells were cultured in mTeSR medium supplemented with zFGF and 1 μg/mL puromycin (Sigma-Aldrich, P9620) for 48 hours to enrich for edited cells. Surviving colonies were expanded under standard mTeSR plus zFGF culture conditions. Genomic DNA was isolated and the targeted region amplified using primers 5′-GCTCTTGGAAACCAGGGCCACC-3′ and 5′-TCAGGAGAACCACCACGGGAACT-3′. Editing outcomes were confirmed by Sanger sequencing and analyzed using the Inference of CRISPR Edits (ICE) algorithm. Multiple independent CHAMP1 KO clones carrying biallelic frameshift-inducing deletions in exon 3 were identified, resulting in premature termination codons. Cells were routinely tested and confirmed negative for mycoplasma contamination.

### Generation of iPSC-derived NPCs

All iPSCs were differentiated into Neural progenitor cells (NPCs) using the STEMdiff SMADi Neural Induction kit (STEMCELL Technologies, #08581). Embryoid Body (EB) formation was completed by plating 2 million cells per well of an AggreWell 800 24-well plate (STEMCELL Technologies, #34811) coated with Anti-Adherence Solution (STEMCELL Technologies, #07010). After five days, the EBs were plated onto one well of a Geltrex (Gibco, #A1413202) -coated 6-well plate. Rosette quality was assessed every day until day 12; only plates with <75% neural rosettes moved forward to the selection and replating stage. Neural rosettes were selected using STEMDIFF Neural Rosette selection media (STEMCELL Technologies, #05832) and replated onto one well of a Geltrex (Gibco, #A1413202) - coated 6-well plate. Full-medium changes with STEMdiff SMADi Neural Induction kit medium were continued until 4-6 days post-plating, until the cells could be passaged. NPC stocks were expanded using STEMdiff Neural Progenitor Medium (STEMCELL Technologies, 05833) on 6-well plates and cryopreserved using STEMdiff Neural Progenitor Freezing Medium (STEMCELL Technologies, #05838). NPC identity was validated using immunocytochemistry and fluorescent microscopy of NPC markers such as PAX6(Abcam, #Ab195045), Nestin(Abcam, #Ab22035), and SOX2(ThermoFisher Scientific, #MA1-014).

### Generation of iPSC-derived Glutamatergic Neurons

NPCs were plated on a poly-L-ornithine (Sigma, #P4957) and laminin (Sigma, #L2020) -coated 24-well plates. NPCs were plated at 100,000 cells per well of a 24-well plate and differentiated into forebrain neuron precursor cells by feeding 0.5mL STEMdiff Forebrain Neuron Differentiation kit (STEMCELL technologies, #08600) daily for five days. The generated neuronal precursors were fed half media changes every other day using the STEMdiff Forebrain Neuron Maturation kit (STEMCELL technologies, #08605) until maturation. Days in vitro (DIV) zero in this paper is used to delineate this change to STEMdiff Forebrain Neuron Maturation media. Neuronal identity was validated using immunohistochemistry and fluorescent microscopy of neuronal maturation markers such as MAP2(Abcam, #Ab183830), NEUN (Abcam, #Ab177487), and VGLUT (ThermoFisher Scientific, MA5-31373).

### Immunocytochemistry

All cells on coverslips in 24-well plates were fixed in 0.5mL of 4% PFA for 10 minutes and washed three times in 1X PBS prior to staining. PBS was then removed and replaced with 0.5mL of PBS with 0.25% Triton X-100 (ThermoFisher Scientific, #A16046.AE) for 10 minutes. Coverslips were washed with 1x PBS for 5 minutes and then incubated in 0.5mL of 1x PBS with 10% goat serum (Gibco, #16210064) for 1 hour at room temperature. Coverslips were then incubated overnight at 4°C with 0.5mL of primary antibody solution which contains 0.2% Triton X-100 in 1x PBS, 2% goat serum, and 1:1000 of the primary antibody. Primary antibody solution was removed and washed three times with 0.5mL of 1x PBS for five minutes per wash. 0.5mL of secondary antibody mix with 2% goat serum, 1:1000 of the secondary antibody in 1X PBS was then added to the coverslips for 1 hour at room temperature. Coverslips were then washed with 1x PBS for a total of three washes and mounted using Flouromount-G w/ DAPI (ThermoFisher Scientific, #00-4959-52). Coverslips were maintained in the dark at room temperature until imaging, then stored at 4°C. All antibodies used for immunocytochemistry are listed in Table 2. Imaging was performed using the STELLARIS Confocal Microscope platform and LAS X software (Version: 4.8.0.28989) at 20x, 40x, or 63x magnification.

### Mitotic Arrest and Analysis

NPCs were plated at 75,000 cells per coverslip (ThermoFisher Scientific, #NC1513283) in one well of a 24-well plate. 24-48 hours after plating, when plate confluency reached around 70%, cells were incubated with 100 ng/mL of Nocodazole (ThermoFisher Scientific, #12-281-0) in STEMdiff Neural Progenitor Medium (STEMCELL Technologies, #05833) for 16 hours^30^. After briefly washing three times in PBS, cells were incubated for 1 hour in maintenance media, then fixed in 4% PFA for 10 minutes, and stored at 4°C until staining. The coverslips were stained with anti-CENP-A (Cell Signaling Technology, #2186S) and anti-γ-tublin (Sigma-Aldrich, t6557-100ul) at a 1:1000 concentration and were mounted using Flouromount-G w/ DAPI (ThermoFisher Scientific, #00-4959-52). The secondary antibodies used were Alexa Fluor 488 goat anti-mouse (ThermoFisher Scientific, #A11001) and Alexa Fluor 680 goat anti-rabbit (ThermoFisher Scientific, #A-21077) at a 1:1000 concentration. Slides were imaged using the STELLARIS Confocal Microscope platform and LAS X software (Version: 4.8.0.28989) at 40x magnification. Ten images were taken per cell line over three coverslips, with a total of three biological replicates per cell line. Cell cycle state was determined by manually counting using Fiji (Version: 1.8.0_345).

### Cellular Proliferation Assay

Neural progenitor cells were seeded at 50,000 cells per well in 24-well plates. Cells were plated and fixed at three passage numbers (5, 10, and 15) with 12 wells per passage per line. Cells were fixed with 4% paraformaldehyde at 24 (0 hour timepoint), 48 (24 hour timepoint), and 72 (48 hour timepoint) hours and stained with 1µg/mL of DAPI (Novus Biologicals, #NBP2-31156-1mg). Cells were counted using an ImageXpress Pico Automated Cell Imaging System and CellReporterXpress analysis program (version 2.9.4.19394). 100% of the well was acquired using the associated IXP-38-570-2141 scope at the 4x magnification. The cell count analysis program was selected with DAPI as the fluorescent channel.

Proliferation was assessed at 24 and 48 hours across three passage numbers (5, 10, and 15) with 12 samples per passage per cell line. Cell counts were log₂-transformed prior to analysis. Doubling time was calculated as: Td = (t₂ − t₁) × ln(2) / (ln(N₂) − ln(N₁)), where t₁ = 24 h, t₂ = 48 h, and N₁, N₂ are the mean cell counts at the respective timepoints. Pairwise differences among cell lines were assessed using a linear mixed effects model (log₂ cell count ∼ Cell Line + Time + (1 | Passage)), with passage modeled as a random intercept to account for repeated measures across passages. Estimated marginal means and pairwise contrasts were computed in R using the emmeans package with Tukey correction for multiple comparisons. Statistical significance was defined as p ≤ 0.05 and denoted as: ns (p > 0.05), * (p ≤ 0.05), ** (p ≤ 0.01), *** (p ≤ 0.001).

### Camptothecin (CPT) DNA Damage Assay

NPCs were treated with 1 µM CPT in Neural Progenitor medium for 1 hour, washed with PBS and the media was replaced. Following incubation at 37°C for 4 hours, cells were fixed in 4% paraformaldehyde. Immunostaining was performed using anti-CyclinA (Sigma-Aldrich, #C4710), anti-Nestin (ThermoFisher Scientific, #PA5-47378), and anti-pRPA2 (ThermoFisher Scientific, #501557195) primary antibodies at a 1:1000 concentration. The secondary antibodies used were Alexa Fluor 546 goat anti-chicken (ThermoFisher Scientific, #A-11040), Alexa Fluor 488 goat anti-mouse (ThermoFisher Scientific, #A11001) and Alexa Fluor 680 goat anti-rabbit (ThermoFisher Scientific, #A-21077) at a 1:1000 concentration. Coverslips were mounted using Flouromount-G w/ DAPI (ThermoFisher Scientific, #00-4959-52). The slides were imaged using the STELLARIS confocal microscope platform and LAS X software (Version: 4.8.0.28989) at 63x magnification. Foci were quantified using the Fiji macro “Foci Analyzer” in cells positive for both Nestin and CyclinA. Three coverslips were stained and quantified per round of each line, with a total of three biological replicates per cell line. All NPCs were treated and fixed through passages 3, 4, and 5.

Foci counts per cell were [log(x + 1)] prior to analysis and only cells with foci counts greater than one were included. Differences in foci count across cell lines were assessed using linear mixed-effects models (LME) fitted with the lme4 and lmerTest packages in R. The model was specified as Foci ∼ line + (1 | rd), where line was the fixed effect of interest and (1 | rd) was a random intercept for an experimental round. When the random-effect variance was estimated at or near zero (singular fit), the model was simplified to an ordinary linear model (Foci ∼ line), as round-to-round variation was considered negligible for the marker. Estimated marginal means (EMMs) were computed for each cell line using the emmeans R package, and all pairwise contrasts between lines were derived from these adjusted means with p-values corrected for multiple comparisons using the Tukey method. Standardized effect sizes (d) were calculated as the estimated pairwise difference divided by the residual standard deviation of the model (small: 0.2, medium: 0.5, large: 0.8). Results are reported as the estimated coefficient (β) with 95% confidence intervals, d-value, and p-value. Statistical significance was defined as p ≤ 0.05 and is denoted as follows: * (p ≤ 0.05), ** (p ≤ 0.01), *** (p ≤ 0.001). Non-significant comparisons are indicated as ns.

### Neuronal Morphology and Neurite Length Analysis

Coverslips of Forebrain neurons ranging from DIV 30-40 and DIV 60-70 were fixed at these timepoints with 4% PFA for 10 minutes and stained with the primary antibodies anti-PAX6 (Abcam, #ab195045) and anti-MAP2 (Abcam, #ab183830) at a 1:1000 concentration to visualize both NPC and neuronal populations within the culture. The secondary antibodies used were Alexa Fluor 488 goat anti-mouse (ThermoFisher Scientific, #A11001) and Alexa Fluor 680 goat anti-rabbit (ThermoFisher Scientific, #A-21077) at a 1:1000 concentration. Coverslips were mounted using Flouromount-G w/ DAPI (ThermoFisher Scientific, #00-4959-52).The slides were imaged using the STELLARIS Confocal Microscope platform and LAS X software (Version: 4.8.0.28989) at the 40x magnification. Ten images were taken between three coverslips, with a total of three biological replicates and 30 images per cell line and timepoint. Only MAP2 positive cells within these images were analyzed using IMARIS version 11.

### Whole-cell patch-clamp electrophysiology and data analysis

Neurons were recorded at two maturation timepoints: DIV 30-40 and DIV 60-70. NPCs were seeded on round glass coverslips (Fisher Scientific, #NC1513283) at 100,000 cells per coverslip and were differentiated into glutamatergic neurons following the protocol above. On recording day, coverslips were transferred from their original culture dish using fine-tipped forceps (Fisher Scientific, #NC1724507) and perfused on the electrophysiology rig with artificial cerebrospinal fluid (pH 7.4, 126 mM NaCl, 2.5 mM KCl, 2 mM CaCl_2_, 26 mM NaHCO_3_, 1.25 mM NaH_2_PO_4_ and 10 mM glucose) and bubbled at RT (25 ± 1°C) with 95% O_2_ and 5% Co_2_. Neurons were visualized under the 40X water objective on an Olympus microscope and recorded using a MultiClamp 700B amplifier. Borosilicate glass microelectrodes between 4-6 Ωm were pulled with pipette puller and filled with potassium-gluconate internal solution containing biocytin for post-hoc labeling (pH 7.3, 290-310 mOsm, 126 mM K-gluconate, 4 mM KCL, 0.3 mM Na_2_-ATP, 4 mM Mg-GTP, 10 mM phosphocreatine, 10 mM HEPES, X % biocytin). Recordings were low pass filtered at 3 kHz and digitized at 20 kHz (Axon Digidata 1550B, Molecular Devices) and data was collected and analyzed using pClamp10.7 and Clampex 10.7. Statistical analysis was conducted using Graphpad Prism 11.0.2. Input resistance was measured in current-clamp mode as the slope of linear fits of current-voltage plots using 1s current injection steps (−10 pA). Series resistance was monitored throughout the recording, and cells exhibiting a change in series resistance ≥ 20% were excluded from all analysis.

We used the following recording programs:

1. Current clamp steps (0 pA for 100ms, steps from −60 pA to +120 pA, 20 pA each, for 1s) to verify the ability of cells to generate action potentials in response to depolarization current steps.
2. Voltage clamp steps (from −90mV to +20 mV in 10mV increments and 1s duration) from a starting potential of −70mV to investigate the presence and amplitude of Na^+^ and K^+^ ionic currents.

### Single-Nuclei RNA-seq sample preparation

One well of a 24-well from three different batches of neurons aged DIV 30 and DIV 60 were frozen down prior to processing. This nuclei preparation protocol was adapted from the 10x protocol CG000124. The cell vials were thawed until about 25% of the sample remained frozen. The sample was doused with 1mL of ACCUTASE (Stemcell Technologies, #07920) and incubated at 37°C for 5 minutes (DIV30) or 10 minutes (DIV60). The sample was quenched with DMEM/F-12 (Gibco, #11320082), centrifuged at 400g and 4°C for 5 minutes. Supernatant was removed and the cell pellet was resuspended in 200µl of Lysis buffer for 15 minutes. Lysis buffer contains 2µL Tris-HCl (ThermoFisher Scientific, #15567-027), 0.4µL NaCl (ThermoFisher Scientific, #AM9760G), 0.6µL MgCl_2_ (ThermoFisher Scientific, #AM9530G), 2µL 10% IGEPAL (Sigma-Aldrich #I8896-50ml), and 195µL 1X PBS. 800 µl of Nuclei wash buffer was used to quench lysis. Nuclei wash buffer contains (300µL 10% bsa in PBS (ThermoFisher Scientific, #37525), 2.685 mL 1x PBS, 15µL of Rnase inhibitor (NxGen, #30281-1). Samples were triturated with a glass pipette until there were no visible clumps. The samples were then centrifuged at 500g and 4°c for 5 minutes, supernatant removed until there was 100ul left in the tube and was resuspended in 900µl of Nuclei wash buffer. The samples were filtered using a 40µm strainer (Fisherbrand, #22363547), centrifuged at 500g and 4°c for 5 minutes. The supernatant was then removed, and the pellet was resuspended in 100ul of Nuclei wash buffer. The nuclei were counted using AOPI (Revity, #CS2-0106-5ml) and the Revity Cellometer Ascend following the manufacturer instructions. The nuclei were processed using the Chromium Single Cell Universal 3’ v2 Gem-x kit following the 10x genomics protocol CG000731. 20K cells/sample were loaded onto the chip and processed by a Chromium X series. Libraries were sequenced by Azenta Life Sciences.

### Single-Nuclei RNA-seq data processing

Raw single nuclei sequencing data were processed with Cell Ranger (v6.1.2)^31^. Cellranger mkfastq command was used to demultiplex the different samples and cellranger count command was used to generate gene-nuclei expression matrices. Sequences were aligned to the GRChg38 reference genome; ambient RNA contamination was inferred and removed using CellBender (v0.232) with standard parameters. Downstream analysis was performed in R with Seurat (v4.1.0)^32^ and customized R scripts. Data from different samples (two batches per stem cell line) were merged into a unique single nuclei object. We applied the following QC filters to retain high quality nuclei: > 200 expressed genes, < 25000 UMIs, and < 5% mitochondrial transcripts. Genes located in the mitochondrial genome were removed. Doublets were removed using scDblFinder (v1.8.0)^33^ for a resultant expression matrix with 17333 genes and 140610 nuclei. The sctransform normalization method in Seurat was used to correct technical variance in read depth and gene detection in individual iPSC lines. To regress out mitochondrial gene percentage and sequencing depth, we used var.to.regress in sctransform. To control for inter-donor variability and other technical effects, samples were then integrated across all the 4 iPSC lines and time points using reciprocal PCA projection method with 3000 variable features (genes). Dimensionality reduction by PCA was used to minimize technical noise in clustering, and the first 50 PCs were selected for UMAP embedding and clustering. We used Louvain algorithm with a resolution of 0.2 to identified initial clusters. Cluster-marker genes were identified for each cluster using the wilcoxauc function in presto (https://github.com/immunogenomics/presto).

### Cellular potency and developmental trajectory

Single-cell developmental potential was estimated using the R package CytoTRACE2 (v1.0.0)^34^, which predicts the differentiation state of individual cells from their transcriptomes and returns both a discrete potency category (differentiated, unipotent, oligopotent, multipotent, pluripotent) and a continuous potency score (range 0–1); predicted values are smoothed by diffusion and rescaled through a combination of binning and k-nearest-neighbor smoothing, with scores closer to 1 indicating a less differentiated state with higher developmental potential and scores closer to 0 a more differentiated state. To enable unbiased cross-genotype comparison, CytoTRACE2 was run on raw counts separately for each line: the control line (CHAMP1^+/+^) was scored first to define a reference potency distribution, and each mutant line (iCHAMP1^-/-^, CHAMP1A, CHAMP1B) was then scored independently. A differentiation pseudotime was defined as 1 − potency score and rescaled to 0–100 and mutant pseudotime values were quantile-normalized (anchored) to the control distribution so that all lines were placed on a common developmental axis. Potency categories and scores were projected onto the UMAP embedding to visualize developmental state across cell types and genotypes.

### Cellular composition analysis

Changes in cellular composition were assessed using sccomp (v2.1.30)^35^, which implements a Bayesian framework based on sum-constrained beta-binomial distributions. We applied the following model: ∼ condition + replicate + (condition | timepoint) + (condition | replicate). Cellular composition was evaluated both globally and at each time point separately. Differential abundance results were further validated using scProportionTest (v0.0.0.9)^36^.

### Differential expression analysis

Genes differentially expressed per cell type and genotype were identified using MAST (v1.32.0)^37^, implemented through Seurat’s FindMarkers function (assay = “RNA”). Within each annotated cell type, each CHAMP1-mutant line (CHAMP1A, CHAMP1B, and iCHAMP1^-/-^) was compared against the control line (CHAMP1^+/+^) as the reference, pooling both differentiation time points. The percentage of mitochondrial transcripts (pMito), cell-cycle phase, and time point (DIV 30/DIV 60) were included as latent variables to account for technical and biological covariates, and only genes detected in at least 30% of cells in either group (min.pct = 0.3) were tested. *P* values were corrected for multiple comparisons using the Benjamini-Hochberg false discovery rate (FDR), and genes were considered significantly up- or down-regulated at an FDR-corrected P value < 0.05 and an absolute log2 fold-change > 0.5. The resulting DEG sets were used as input for the downstream functional and disease enrichment analyses.

### Functional Enrichment

For functional enrichment of the differentially expressed genes, we used scToppR package (v0.99.4)^38^, an R interface to the ToppGene/ToppFun functional enrichment suit, requiring between 5 and 5,000 genes per term and returning up to 50 enriched terms. Enrichment was assessed across Gene Ontology Biological Process, Gene Ontology Molecular Function, and Disease categories, with P values corrected using the Benjamini-Hochberg (BH) method. For visualization, the top five terms per category and cell type were displayed as the −log10 of the BH-adjusted P value. Furthermore, to identify synaptic processes potentially affected in mature neurons, we performed Gene Ontology (GO) enrichment analysis using SynGO. Differentially expressed genes (DEGs) were submitted to the SynGO portal (https://www.syngoportal.org), from which GO annotations and corresponding sunburst visualizations were generated.

### GWAS-risk loci and Gene Set enrichment

To evaluate the association of individual cells with neurodevelopmental and psychiatric disorders, we applied scDRS (v1.0.2)^39^ to our single-nucleus dataset. For each disorder (ASD, ADHD, and epilepsy), putative risk genes and their gene-level association statistics were derived from GWAS summary statistics^40–42^ using multi-marker analysis of genomic annotation (MAGMA)^43^. Each putative disease gene was weighted by its MAGMA z-score from GWAS and inversely weighted by its gene-specific technical noise level estimated within our single-nucleus data, generating cell-specific raw disease scores. For downstream analysis, we performed cell-type-level assessments to link broad cell types to each disorder and to examine heterogeneity in disease association across cells within each broad cell type, using the scDRS perform_downstream() function with default settings. To address multiple testing, the FDR was calculated using the Benjamini–Hochberg method. For gene set enrichment analysis, we used the SFARI ASD genes (https://gene.sfari.org)^44^. Enrichment was assessed using Fisher’s exact test in R, specifying an alternative hypothesis of “greater” and a confidence level of 0.99. Odds ratios (ORs) were calculated, and multiple testing correction was performed using the Benjamini– Hochberg false discovery rate (FDR) method.

### Coexpression Network analysis

High-dimensional weighted gene co-expression network analysis was performed using the hdWGCNA package (v0.4.03)^45^, a systems-level approach that uses unsupervised learning to identify groups of co-expressed genes (modules) with differential activity across conditions. As our dataset comprised four annotated cell types, an independent signed co-expression network was constructed for each cell type rather than collapsing clusters into broader groups. For each network, genes expressed in at least 5% of cells within the corresponding cell type were retained (gene_select = “fraction”). To counter the gene dropout inherent to snRNA-seq, metacells were aggregated within each genotype using MetacellsByGroups on RNA-based UMAP coordinates with k = 25 nearest neighbors, a maximum of 10 cells shared between metacells, and a minimum of 30 cells per group, after which metacell expression was normalized. Expression matrices for network construction were assembled across all four lines (CHAMP1^+/+^, iCHAMP1^-/-^, CHAMP1A, CHAMP1B). Soft-power thresholds were tested with TestSoftPowers using a signed network and the biweight midcorrelation (bicor), and the lowest soft power yielding an appropriate scale-free topology fit was selected for each cell type (NPC, 8; ccNPC, 4; immature neurons, 5; mature neurons, 6). Signed networks were constructed with ConstructNetwork using bicor, deepSplit = 4, and a merge cut height of 0.10. Summarized module expression values (module eigengenes, MEs) were calculated with ModuleEigengenes and corrected across donors with Harmony, and eigengene-based connectivity (kME) was calculated for each gene with ModuleConnectivity; the grey (unassigned) module was excluded from all downstream analyses. Modules were renamed by cell type (e.g., MatureNeurons-M4), and per-cell module activity scores were derived from the top 25 hub genes of each module using the UCell method (ModuleExprScore) (v2.10.1)^46^. Module hub-gene networks were visualized using the top 30 hub genes and top 150 connections per module. To test for genotype-associated changes in module activity, MEs were modeled as a function of genotype (ME ∼ condition) with CHAMP1^+/+^ as the reference, and *P* values were corrected across modules using the Benjamini-Hochberg procedure (FDR; *FDR < 0.05, **FDR < 0.01, ***FDR < 0.001). Module activity was further related to developmental progression by correlating MEs with CytoTRACE2 potency and scaled pseudotime (Spearman’s rho, BH-corrected) and by plotting binned, min-max–normalized eigengene activity along pseudotime for each genotype. Each module was functionally annotated by Gene Ontology (Biological Process) over-representation analysis using clusterProfiler (v4.14.6)^47^, displaying the top five terms per module.

### Statistical Analysis

The statistical analyses in this study were conducted using Graphpad Prism (11.0.2) or R using the emmeans package. Graphs denote the means of cellular and neuronal properties with Standard Error of the Mean (SEM) as the error bars. Unless otherwise stated, one-way or two-way analysis of variance (ANOVA) with multiple comparisons and Tukey’s post-hoc multiple comparisons test were conducted between mutant cell lines (CHAMP1A, CHAMP1B, and iCHAMP1^-/-^) compared with a control line (CHAMP1^+/+^) within the same timepoint or treatment group. Statistical significance as a P value <0.05 is noted within figures or tables, as *p<0.05, **p<0.01, ***p<0.005, and ****p<0.001. Comparisons without these notations are considered statistically not significant.

## RESULTS

### CHAMP1 Patient-derived Neural Progenitor cells exhibit HR deficiency and aberrant proliferation

CHAMP1 promotes homologous recombination (HR)-mediated repair of DNA double-strand breaks (DSBs) by facilitating DNA end resection^17^. Consistent with prior findings demonstrating that CHAMP1 loss-of-function impairs HR in patient-derived fibroblasts following camptothecin (CPT) treatment^48^, we investigated whether neural progenitor cells (NPCs) harboring heterozygous CHAMP1 mutations exhibit similar defects in HR-mediated DSB repair (Fig. 2A; Methods). In this acute DNA damage model, CHAMP1B mutant NPCs showed significantly reduced pRPA2 foci following CPT treatment compared to CHAMP1^+/+^ (p = 0.00411; ANOVA with Tukey correction, Fig. 2B-C; Table S2), consistent with impaired DNA end resection. CHAMP1A exhibited the most pronounced HR deficiency, with markedly fewer pRPA2 foci than CHAMP1^+/+^ (p < 0.0001; ANOVA with Tukey correction, Fig. 2B-C). Notably, iCHAMP1^−/−^ cells showed no significant reduction in pRPA2 foci compared to CHAMP1^+/+^ (p = 0.992, ns), suggesting that complete CHAMP1 loss engages compensatory repair mechanisms distinct from those disrupted by heterozygous mutations (Fig. 2B-C). These mutation-specific alterations in DNA end resection suggest that CHAMP1 regulates DSB repair in an allele-dependent manner and highlight heterogeneity in repair phenotypes across CHAMP1 mutation classes. To determine the functional consequences of these defects, we assessed NPC proliferation by quantifying cell number at 24 and 48 hours post-plating. CHAMP1^+/+^ cells exhibited the shortest doubling time (34.4 hours), followed by CHAMP1A (35.0 hours), CHAMP1B (43.7 hours), and iCHAMP1^−/−^ (51.1 hours), indicating progressively reduced proliferative capacity across mutant genotypes. These findings were corroborated using an independent neurotypical control line (CHAMP1_2^+/+^), which recapitulated the proliferative advantage of CHAMP1^+/+^ cells over all mutant genotypes, confirming that the observed differences reflect genotype-specific effects rather than clonal variability (Fig. S2). Pairwise comparisons using estimated marginal means demonstrated that CHAMP1^+/+^ cells proliferated significantly more than all mutant lines (all p < 0.001; ANOVA with Tukey correction). Among mutants, iCHAMP1^−/−^ cells exhibited significantly reduced proliferation compared to both CHAMP1A (p = 0.007) and CHAMP1B (p = 0.024), while no significant difference was observed between CHAMP1A and CHAMP1B (p = 0.975), indicating that CHAMP1 dosage critically regulates NPC proliferative capacity.

**Figure 1.**
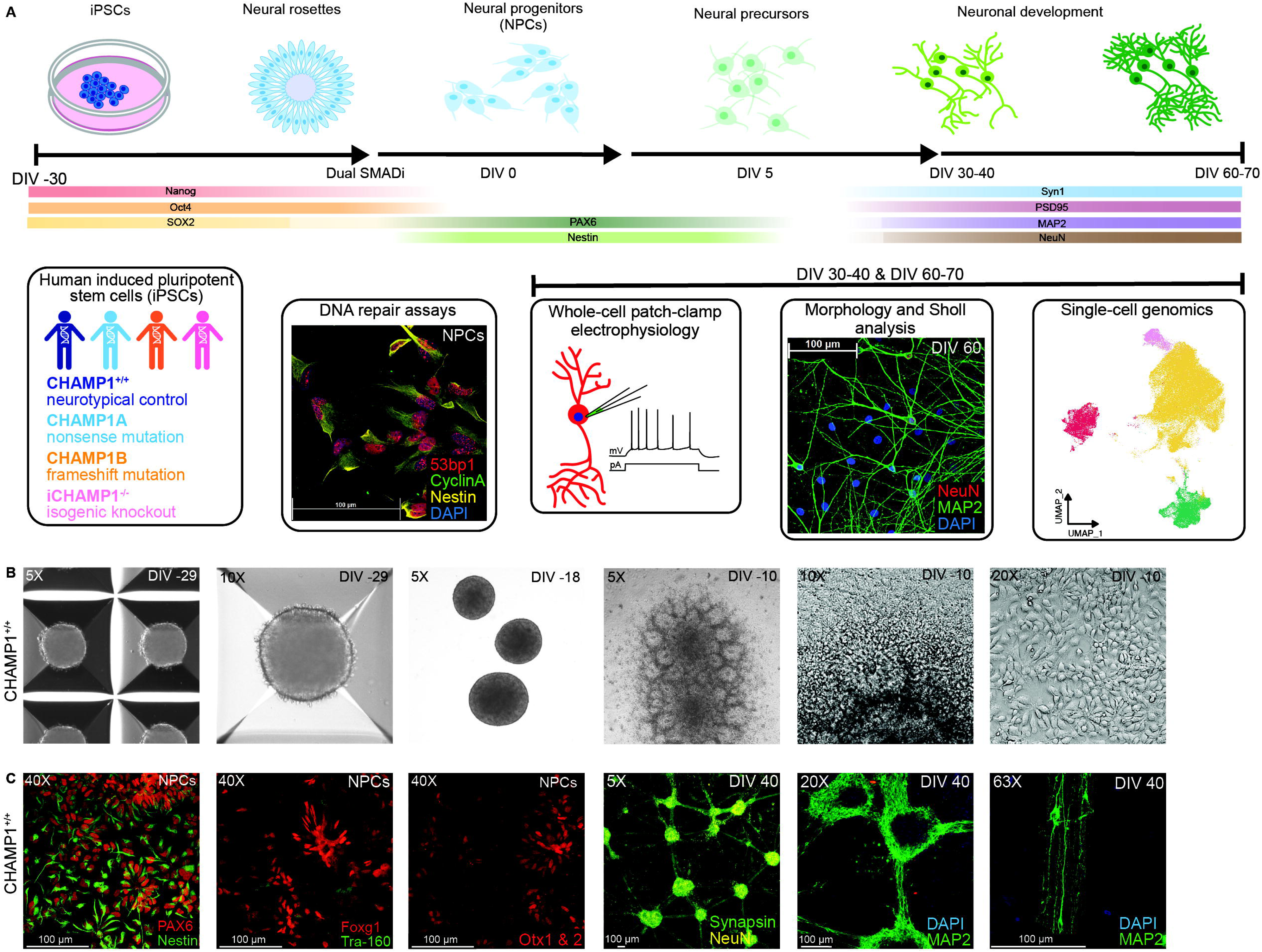
Modeling neurodevelopment in patient-derived iPSC based neuronal cultures. ***A***, Timeline of maturing patient-derived iPSC lines into neurons using Dual SMADi protocol. This study included 4 cell lines; one neurotypical control, two CHAMP1 patient mutations, and one isogenic knockout. DNA repair assays were conduced in NPCs. Experiments involving whole-cell patch-clamp electrophysiology, post-hoc morphology and Sholl analysis, and Single-cell genomics were conducted at DIV 30-40 and DIV 60-70 in maturing neurons. Image was created with BioRender.com. ***B***, Example brightfield images of iPSC to NPC protocol in CHAMP1^+/+^ cells. One day after centrifuging iPSCs into one well of an Aggrewell 800 plate, Day −29 embryoid bodies (EBs) form. EBs continue to grow in suspension. EBs are replated onto geltrex coated plates and neural rosettes form. ***C***, Confocal images of CHAMP1^+/+^ cells before and after neural rosette selection when NPCs are isolated. NPCs express markers of neural progenitor cells including PAX6, Nestin, Foxg1, and Otx 1&2, with low expression prevalence of stem cell marker Tra-160. Neurons at DIV 40 express markers of mature neurons including Synapsin, NeuN, and MAP2.

**Figure 2.**
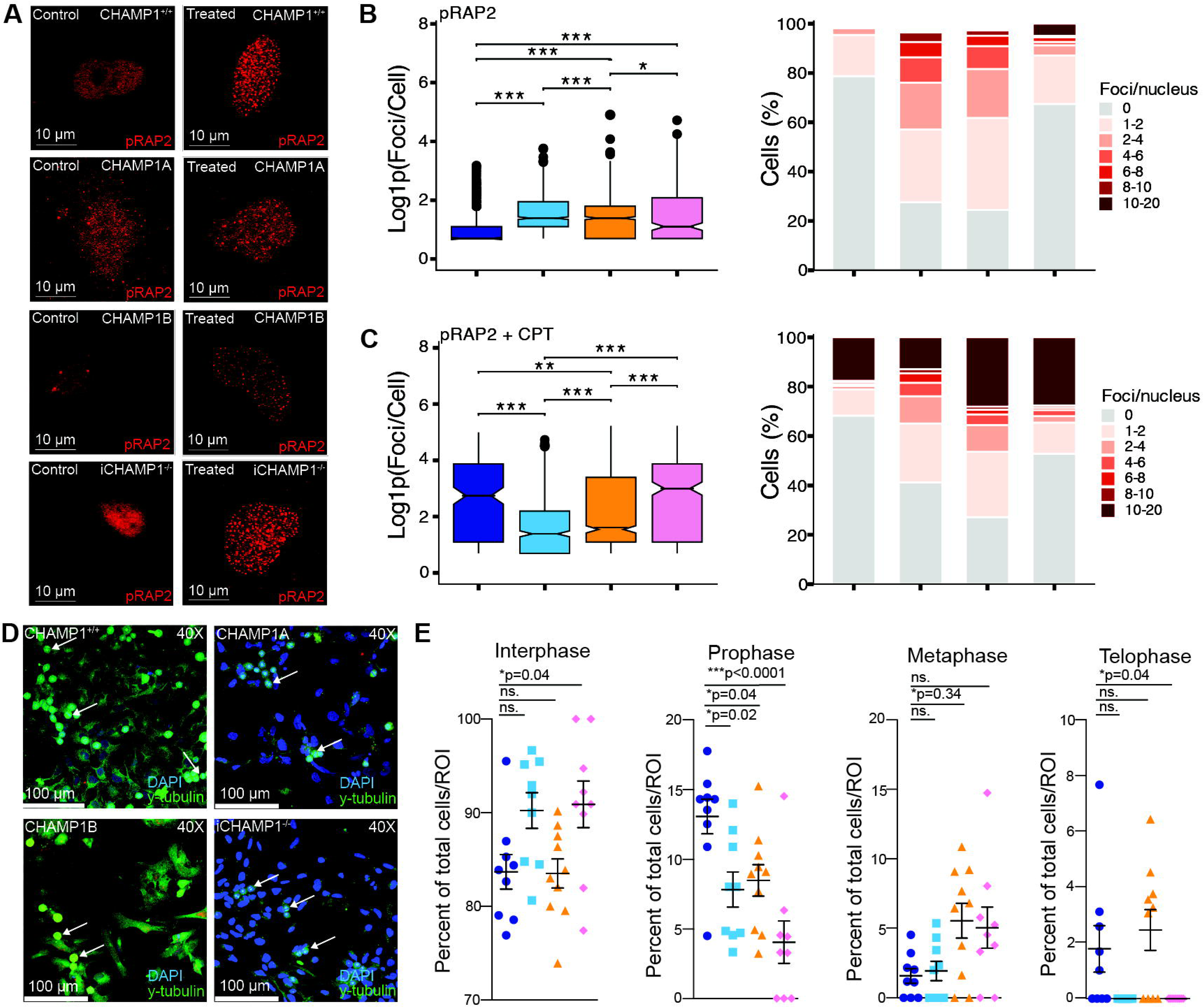
Patient-derived NPCs exhibit HR deficiency and aberrant mitotic progression. ***A,*** Representative confocal images of pRPA2 immunostaining in CHAMP1⁺^/^⁺, CHAMP1A, CHAMP1B, and iCHAMP1^⁻/⁻^ NPCs under basal (Control) and DNA damage (Treated) conditions. NPCs were exposed to 1 µM CPT for 1 hour, allowed to recover for 4 hours, fixed, and immunostained with anti-pRPA2 antibodies. Scale bar: 10 µm. **B,** Quantification of pRPA2 foci in untreated NPCs. Left: log1p-transformed foci counts per cell shown as box-and-whisker plots; the horizontal bar indicates the median. Right: stacked bar charts showing the distribution of cells across foci-count bins (0, 1–2, 2–4, 4–6, 6–8, 8–10, and 10–20 foci per nucleus), normalized to 100% per line. Only cells with foci counts greater than one were included in quantitative analyses. Data represent >150 cells per line across three biological replicates. ***C,*** Quantification of pRPA2 foci following CPT treatment, displayed as in B. Statistical comparisons are relative to CHAMP1^⁺/⁺^ (ANOVA with Tukey correction; *p<0.05, **p<0.01, ***p<0.001). ***D,*** Representative confocal images of NPCs stained with DAPI (blue) and anti-γ-tubulin (green) to resolve individual mitotic stages. Scale bar: 100 µm. Arrows indicate representative mitotic cells*. **E,*** Quantification of the proportion of NPCs in each mitotic phase (interphase, prophase, metaphase, and telophase) across cell lines, expressed as a percentage of total cells per ROI. Ten images were acquired per line across three biological replicates. Statistical comparisons shown above each phase (ns, not significant; *p<0.05; ***p<0.0001; ANOVA with Tukey correction). Data are shown as mean ± SEM.

Given prior reports linking CHAMP1 depletion to defects in mitotic progression and chromosome segregation^19^, we next examined whether similar abnormalities are present in patient-derived NPCs using γ-tubulin immunostaining to resolve individual mitotic stages. iCHAMP1^−/−^ cells exhibited significant accumulation in interphase relative to CHAMP1^+/+^ (p = 0.04; ANOVA with Tukey correction,. 2D-E), indicative of delayed cell cycle progression, and CHAMP1B cells showed a similar but non-significant trend. Conversely, CHAMP1^+/+^ cells displayed a higher proportion of cells in prophase compared to CHAMP1A (p = 0.02), CHAMP1B (p = 0.04), and iCHAMP1^−/−^ (p < 0.0001), suggesting impaired mitotic entry in CHAMP1-deficient lines (Fig. 2E). While all mutant lines exhibited increased proportions of cells in metaphase, this reached significance only for CHAMP1B (p = 0.034). No significant differences were observed in telophase across genotypes (Fig. 2E).

Collectively, these data establish a mechanistic link between CHAMP1 dysfunction, impaired DNA end resection, and reduced neural progenitor proliferation, providing insight into how allele-specific CHAMP1 mutations may contribute to the neurodevelopmental pathology observed in affected individuals.

### CHAMP1 Mutations Disrupt Membrane Properties of Excitatory Neurons During Neuronal Maturation

To investigate how CHAMP1 mutations affect the maturation and function of cortical excitatory neurons, we generated glutamatergic neuronal cultures from patient-derived and isogenic iPSCs using dual-SMAD inhibition and performed whole-cell patch-clamp recordings at two developmental stages: DIV 30–40, representing early neuronal differentiation, and DIV 60–70, approaching mature excitability. These time points capture key transitions in human iPSC-derived cortical neuron maturation, as spontaneous firing emerges by approximately DIV 20 and intrinsic membrane properties continue to mature through DIV 70^49^.

Across multiple intrinsic membrane properties, including input resistance, access resistance, holding current, and rheobase, we observed consistent, mutation-specific deficits indicative of delayed cellular maturation (Table 1; Table S3). In particular, CHAMP1A, CHAMP1B, and iCHAMP1^-/-^ neurons showed significantly elevated access resistance at DIV 60–70, consistent with poorer space clamp and immature membrane properties, and more depolarized holding currents at one or both time points compared with CHAMP1^+/+^ (DIV 30-40 CHAMP1^+/+^ −20 ± 2.4 pA n=28, CHAMP1A −9.5 ± 0.9 pA n=47, CHAMP1B −11.4 ± 2.6 pA n=31, iCHAMP1^-/-^ −6.7 ± 1.1 pA n=30; DIV 60-70 CHAMP1^+/+^ −17 ± 2.8 pA n=28, CHAMP1A −7.4 ± 2.1 pA n=36, CHAMP1B −4.3 ± 2.7 pA n=34, iCHAMP1^-/-^ −8.3 ± 1.5 pA n=34; ANOVA, mean ± SEM; Table 1). These features are characteristic of electrophysiological immaturity and are commonly reported in iPSC models of neurodevelopmental disorders, suggesting that CHAMP1 mutations disrupt the developmental trajectory of neuronal membrane properties maturation, potentially through altered ion channel trafficking, soma growth, or cytoskeletal organization^50,51^. Together, these findings position CHAMP1 as a critical regulator of human neuronal maturation, linking its roles in genome stability and cytoskeletal function to the establishment of intrinsic excitability. Disruption of these processes likely contributes to circuit-level dysfunction in CHAMP1-associated neurodevelopmental disorders.

### CHAMP1 Mutations Disrupt Waveform Properties at Rheobase and Reduce Action Potential Firing During Early Neurodevelopment *In Vitro*

Because CHAMP1 mutations altered AP waveform maturation at rheobase, we next asked whether these defects extended to repetitive firing, a more stringent readout of intrinsic excitability and functional neuronal maturation^52^. Given that disruption of ASD/NDD-associated regulators can broadly alter firing patterns during development^53,54^, we hypothesized that CHAMP1 mutations would impair repetitive AP generation, potentially reflecting hypoexcitability rather than a seizure-prone phenotype.

Therefore, to assess the role of CHAMP1 in the functional development of neurons in vitro, we performed whole-cell patch-clamp recordings of intrinsic membrane properties and action potential (AP) firing dynamics in iPSC-derived excitatory neurons at DIV 30–40, corresponding to early differentiation, and DIV 60–70, when neurons approach mature excitability. NMDA/AMPA blockers (CPP, DNQX) and the L-type Ca²⁺ channel antagonist nifedipine isolated intrinsic excitability from network inputs (Methods). Compared to CHAMP1^+/+^, CHAMP1 mutations disrupted AP waveform maturation at rheobase, particularly at DIV 30–40, with iCHAMP1^-/-^ neurons showing the most pronounced deficits, including depolarized thresholds, reduced peak amplitudes, depolarized afterhyperpolarizations, and prolonged half-widths (Threshold −25 ± 1.0 mV, Peak amplitude 47 ± 2.9 mV, Anti-peak amplitude −9.5 ± 1.2 mV, Half-width 8.8 ± 0.78 ms; n=22, ANOVA, mean ± SEM; Fig. 3A–F). These are canonical markers of immature excitability and largely resolved by DIV 60–70 (Threshold −30 mV ± 1.1 mV, Peak amplitude 63 ± 4.7 mV, Anti-peak amplitude −7.9 ± 1.4 mV, Half-width 5.1 ± 0.91 ms; n=12, ANOVA, mean ± SEM; Fig. 3A–F; Table S3).

**Figure 3.**
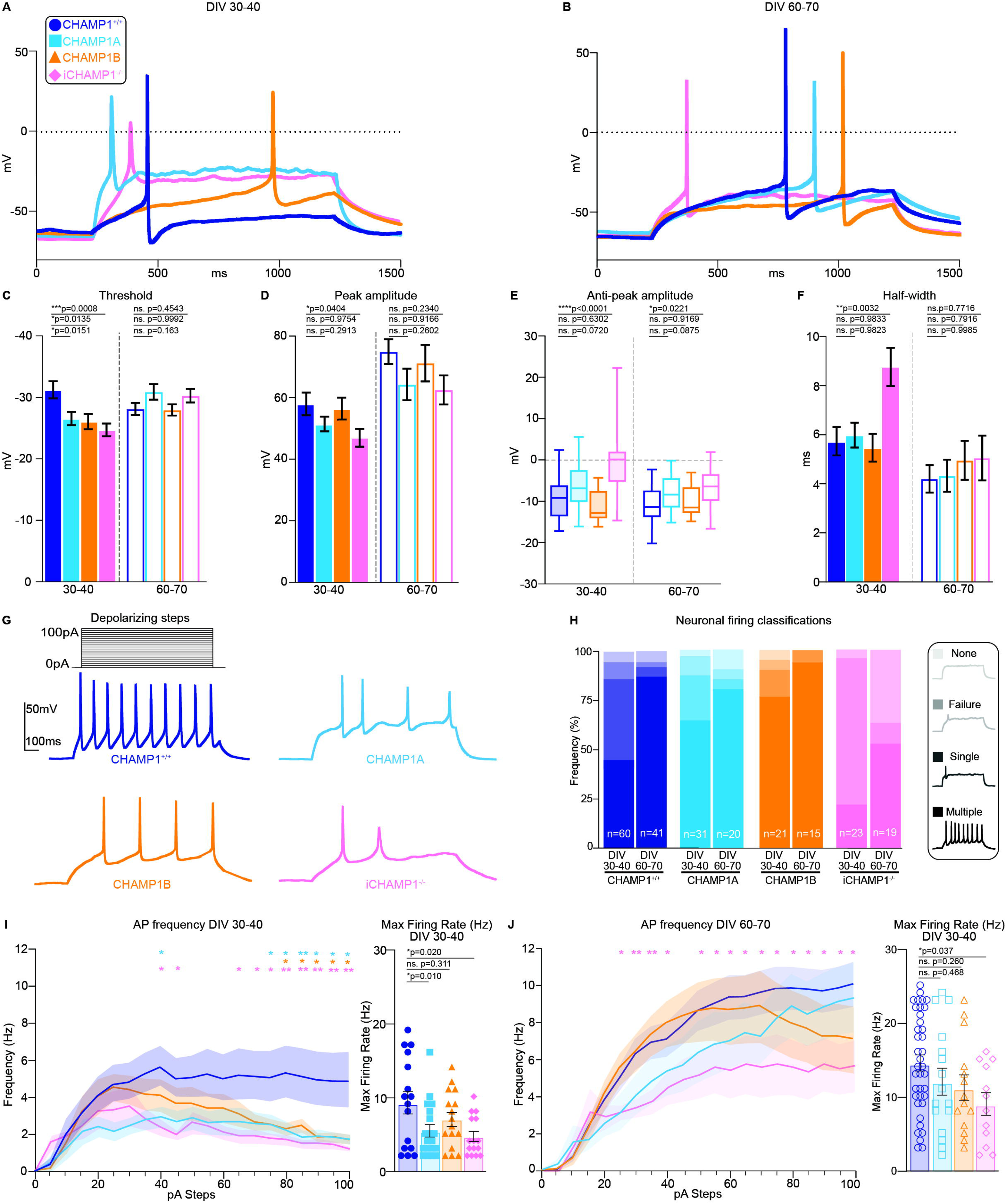
CHAMP1 mutations disrupt waveform properties and action potential frequency. ***A-B***, Example traces of action potentials at rheobase from neurons at DIV 30-40 (a) and DIV 60-70 (b). ***C***, Threshold for AP firing is significantly reduced in magnitude in all groups compared to CHAMP1^+/+^ at DIV 30-40. Threshold is not significantly altered at DIV 60-70. ***D***, Peak amplitude is significantly reduced in iCHAMP1^-/-^ neurons compared to CHAMP1^+/+^ at DIV 30-40 but not DIV 60-70. ***E***, Anti-peak amplitude was reduced in iCHAMP1^-/-^ neurons compared to CHAMP1^+/+^ at both timepoints. ***F***, Half-width was significantly reduced in iCHAMP1^-/-^ neurons compared to CHAMP1^+/+^ neurons at DIV 30-40, but not DIV 60-70. ***G***, 5pA depolarizing steps from 0 to 100pA and representative action potential traces for 4 cell lines. ***H***, Percentage of neurons recorded represented by firing rate profile (None, Failure, Single, & Multiple, lowest to highest color intensity respectively). The percentage of neurons firing multiple action potentials varied across lines and timepoints, but was the lowest in iCHAMP1^-/-^ at both timepoints. ***I***, Action potential frequency at DIV 30-40 and Maximum AP frequency for each cell at DIV 30-40 (left) (within-timepoint statistics, ANOVA, *p<0.05). ***J***, Action potential frequency at DIV 60-70 and Maximum AP frequency for each cell at DIV 60-70 (within-timepoint statistics, ANOVA, *p<0.05). Max AP frequency was significantly decreased in both CHAMP1A and iCHAMP1^-/-^ compared to CHAMP1^+/+^ at DIV 30-40, and in iCHAMP1^-/-^ compared to CHAMP1^+/+^ at DIV 60-70.

Interestingly, we found that action potential threshold was significantly less depolarized across *CHAMP1* mutations compared to CHAMP1^+/+^ at DIV 30-40, whereas no significant difference was observed at DIV 60–70 (DIV 30-40, CHAMP1^+/+^ −31 ± 1.4 mV n=19, CHAMP1A −27 ± 1.1 mV n=27, CHAMP1B −26 ± 1.2 mV n=19, iCHAMP1^-/-^ −25 ± 1.0 mV n=22; DIV 60-70, CHAMP1^+/+^ −28 ± 0.99 mV n=30, CHAMP1A −31 ± 1.3 mV n=17, CHAMP1B −28 ± 0.92 mV n=15, iCHAMP1^-/-^ −28 ± 0.92 mV n=15; ANOVA, mean ± SEM; Fig. 3C). Peak amplitude at rheobase was significantly reduced in iCHAMP1^-/-^ neurons at DIV 30–40 but was not different at DIV 60–70, and it was not significantly altered in CHAMP1A or CHAMP1B at either time point (DIV 30-40, CHAMP1^+/+^ 58 ± 3.7 mV n=19, CHAMP1A 51 ± 2.3 mV n=27, CHAMP1B 56 ± 3.6 mV n=19, iCHAMP1^-/-^ 47 ± 2.9 mV n=22; DIV 60-70, CHAMP1^+/+^ 75 ± 4.0 mV n=35, CHAMP1A 64 ± 5.1 mV n=32, CHAMP1B 71 ± 5.9 n=39, iCHAMP1^-/-^ 63 ± 4.7 mV n=34; ANOVA, mean ± SEM; Fig. 3D). Consistent with this, anti-peak amplitude was significantly more positive in iCHAMP1^-/-^ neurons at both time points (DIV 30-40 −0.39 ± 1.6 mV n=22; DIV 60-70 −7.9 ± 1.4 mV n=12; ANOVA, mean ± SEM; Fig. 3E), indicating impaired depolarization and repolarization during AP firing. Half-width, which typically decreases with neuronal maturation, was also significantly prolonged in iCHAMP1^-/-^ neurons at DIV 30–40 (8.8 ± 0.78 mV n=22; ANOVA, mean ± SEM; Fig. 3F), further supporting an immature electrophysiological state. Together, these findings suggest that CHAMP1 mutations delay the maturation of the cellular machinery required for normal neuronal excitability.

Because CHAMP1 mutations altered AP waveform maturation at rheobase, we next asked whether these defects extended to repetitive firing, a more stringent measure of intrinsic excitability and functional neuronal maturation. As CHAMP1 mutant patient-derived lines could generate action potentials at rheobase, and because CHAMP1A and CHAMP1B showed relatively modest waveform abnormalities, we hypothesized that the main functional consequence of CHAMP1 loss might be impaired repetitive firing capacity rather than complete failure of AP generation. More specifically, given the clinical phenotype of CHAMP1-associated disorder, which is characterized by developmental delay, hypotonia, and neurodevelopmental impairment, we hypothesized that CHAMP1 mutations would produce hypoexcitability rather than hyperexcitability. This prediction is also consistent with prior work showing that ASD- and NDD-associated genes can alter repetitive firing during neuronal maturation^55^.

To test this hypothesis, we quantified repetitive AP firing at 5pA depolarization steps in current clamp mode (Fig. 3G-J; Methods). We evaluated the representation of functional maturation among the neuronal population, both between groups and across time. We classified neurons based on how many APs they fired: none, failure, single, or multiple. Across all groups, the proportion of neurons capable of firing multiple APs increased over time, consistent with progressive maturation in this iPSC-derived cortical neuron model. Interestingly, both CHAMP1A and CHAMP1B neurons had relatively higher representation of the “multiple” categories at DIV 30-40 compared with CHAMP1^+/+^, despite having significantly lower firing rates during depolarizing steps (CHAMP1^+/+^ 40% n=35, CHAMP1A 60% n=31, CHAMP1B 75% n=21, iCHAMP1^-/-^ 22% n=23; Fig. 3H). In contrast, iCHAMP1^-/-^ neuron population was overall less mature at both timepoints, in accordance with the functional readout of reduced repetitive action potential firing (DIV 30-40 22% n=23; DIV 60-70 53% n=19 Fig. 3H). With these results in mind, we quantified the depolarization curves and maximum AP numbers only for neurons that fired 2 or more AP’s during depolarizing current steps (Fig. 3I-J) to normalize the dataset across cell lines and exclude functionally immature neurons (single-firing). At DIV 30–40, all mutant lines showed a marked reduction in AP frequency relative to CHAMP1^+/+^ controls between 80-100 pA steps (Fig. 3I, *p<0.05, **p<0.01, ANOVA, mean ± SEM, left), and pronounced deficits in maximum AP frequency in CHAMP1A and iCHAMP1^-/-^ (Max Firing Rate, CHAMP1^+/+^ 9.2 ± 1.5 Hz n=16, CHAMP1A 5.4 ± 0.84 Hz n=19, CHAMP1B 6.9 ± 0.96 n=16, iCHAMP1^-/-^ 4.6 ± 0.69 Hz n=15; ANOVA, mean ± SEM, Fig. 3I, right), indicating an early deficit in intrinsic excitability that may contribute to later circuit-level dysfunction. At DIV 60–70, AP frequency was no longer significantly different in CHAMP1A or CHAMP1B neurons at either individual pA steps or maximum frequency, whereas iCHAMP1^-/-^ neurons remained significantly reduced relative to CHAMP1^+/+^ controls (Max Firing Rate, CHAMP1^+/+^ 14 ± 1.1 Hz n=38, CHAMP1A 12 ± 1.8 Hz n=16, CHAMP1B 11 ± 1.7 n=14, iCHAMP1^-/-^ 8.9 ± 1.5 Hz n=12; ANOVA, mean ± SEM; Fig. 3J, right). These findings suggest that CHAMP1 is required for the timely acquisition of repetitive firing capacity in developing excitatory neurons.

To validate the metrics used to compare mutant lines with CHAMP1^+/+^ line, we recorded from an additional neurotypical line (CHAMP1_2^+/+^) from a different genetic background at both DIV 30-40 and DIV 60-70 (Fig. S3). We found no significant difference between lines in maximum action potential frequency with increasing current steps at DIV 30-40 or DIV 60-70 (Fig. S3A-B). Interestingly, CHAMP1_2^+/+^ had almost twice as many neurons firing multiple action potentials compared with CHAMP1^+/+^ across the population recorded from at DIV 30-40, but similar proportions at DIV 60-70 (Fig. S3C). While no significant differences were recorded in rheobase threshold or Anti-peak amplitude, there was a slight increase in peak amplitude at DIV 30-40 and a decrease in half-width at both timepoints in CHAMP1_2^+/+^ neurons compared to CHAMP1^+/+^(Fig. S3D-G).

Taken together, these findings demonstrate that CHAMP1 mutations disrupt AP waveform maturation, profoundly impair early repetitive firing capacity, and delay population-level excitability trajectories in developing excitatory neurons.

### CHAMP1 mutations reduce voltage-gated sodium channel current densities

To investigate the underlying mechanisms of the repetitive AP firing deficits observed in CHAMP1 mutants, we examined voltage-gated Na^+^ and K^+^ currents, which serve as critical determinants of membrane excitability (Fig. 4). During AP firing, changes to the membrane voltage activate sodium ion channels and cause a fast inward flux of sodium ions, followed by a slower outward flux of potassium ions. The presence of more functionally active voltage-gated ion channels on the neuronal membrane is directly correlated to AP peak and anti-peak, and can reflect the neuron’s capability for higher AP frequencies^56,57^.

**Figure 4.**
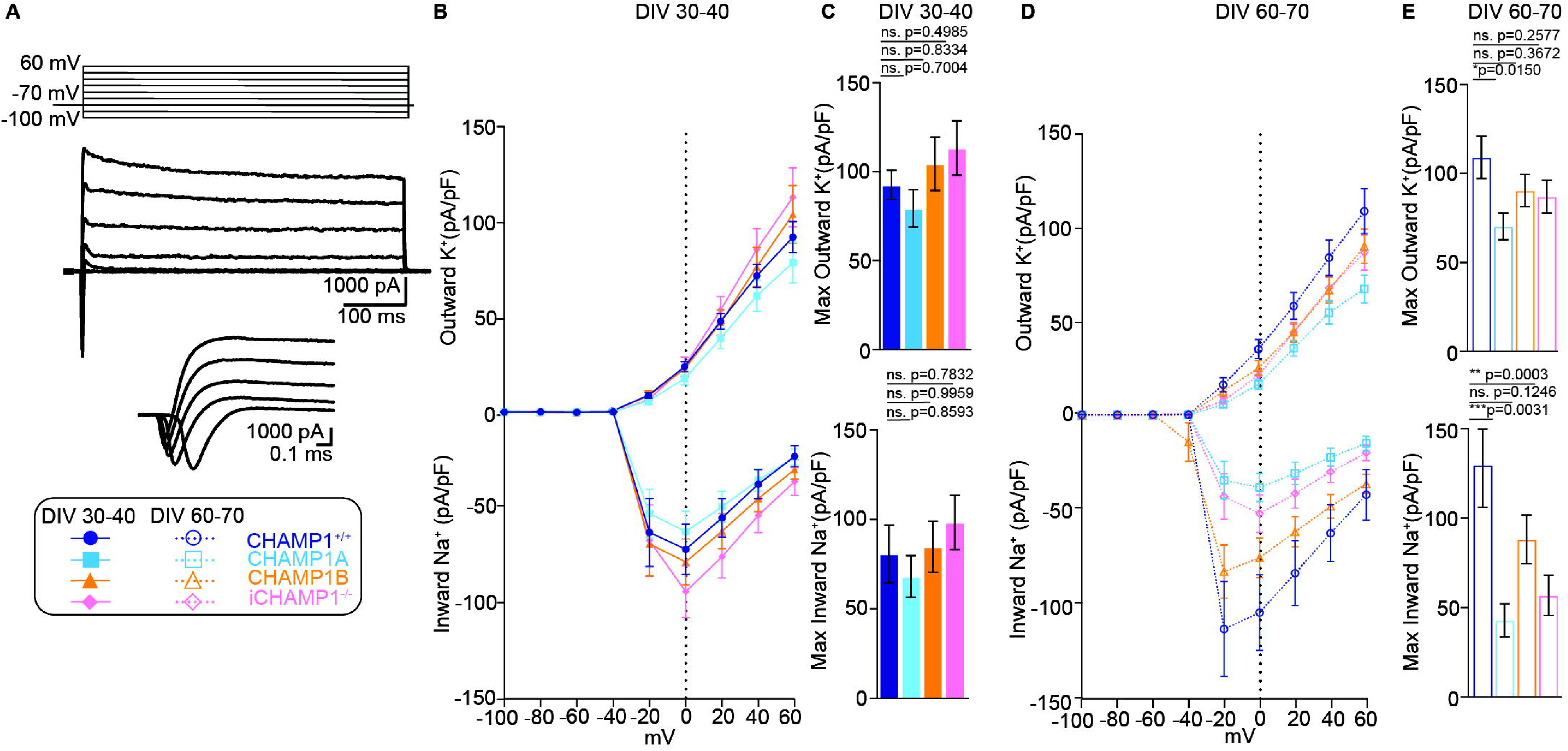
CHAMP1 mutations reduce sodium and potassium current densities. ***A***, Neurons in current clamp receive 20mV steps from −100mV to 60mV to record inward Na+ and outward K+ current density normalized by average cell capacitance within-group at each timepoint (Table 1) (pA/pF). ***B***, Outward K+ (top) and inward Na+ (bottom) current density at DIV 30-40. ***C***, Max outward K+ current density (top) and Max inward Na+ current density (bottom) were unchanged between groups at DIV 30. ***D***, Outward K+ (top) and inward Na+ (bottom) current density at DIV 60-70. ***E***, Max outward K+ current density (top) is significantly reduced in CHAMP1A compared to CHAMP1^+/+^ DIV 60-70. Max inward Na+ current density (bottom) was significantly reduced in both CHAMP1A and iCHAMP1^-/-^ compared to CHAMP1^+/+^ DIV 60-70.

We evaluated the density of inward sodium and outward potassium currents in voltage clamp mode using voltage steps from −100mV to +60mV and reported the normalized current density in pA/pF using the average cell capacitance within-group at each timepoint (Fig. 4A; Table S4; Methods). At DIV 30-40, we did not observe any significant changes in either sodium or potassium currents across groups compared with CHAMP1^+/+^ (Max Outward K^+^; CHAMP1^+/+^ 93 ± 8.2 pA/pF n=25, CHAMP1A 79 ± 11 pA/pF n=23, CHAMP1B 104 ± 15 pA/pF n=14, iCHAMP1^-/-^ 113 ± 15 n=13; ANOVA, mean ± SEM, Fig. 4B-C (top); Max Inward Na^+^; CHAMP1^+/+^ 81 ± 16 pA/pF n=25, CHAMP1A 68 ± 12 pA/pF n=23, CHAMP1B 85 ± 14 pA/pF n=14, iCHAMP1^-/-^ 98 ± 15 pA/pF n=13; ANOVA, mean ± SEM; Fig. 4B-C (bottom)). However, both CHAMP1A and iCHAMP1^-/-^ neurons had significantly reduced inward sodium currents at DIV 60-70 compared to CHAMP1^+/+^, while CHAMP1B neurons were comparable with CHAMP1^+/+^ (Max Outward K^+^; CHAMP1^+/+^ 109 ± 12 pA/pF n=18, CHAMP1A 70 ± 7.5 pA/pF n=22, iCHAMP1^-/-^ 87 ± 9.3 pA/pF n=20; ANOVA, mean ± SEM; Fig. 4D-E (top); Max Inward Na^+^; CHAMP1^+/+^ 103 ± 24 pA/pF n=17, CHAMP1A 43 ± 9.2 pA/pF n=21, CHAMP1B 88 ± 14 pA/pF n=22, iCHAMP1^-/-^ 57 ± 11 pA/pF n=20; ANOVA, mean ± SEM; Fig.4D-E (bottom)). We detected no significant differences in Max Outward K^+^ or Max Inward Na^+^ at DIV 30-40 between CHAMP1^+/+^ and CHAMP1_2^+/+^, no difference in Max Inward Na^+^ at DIV 60-70, but a reduction in Max Outward K+ current at DIV 60-70 in CHAMP1_2^+/+^ (DIV 30-40, Max Outward K^+^; CHAMP1^+/+^ 93 ± 8.2 pA/pF n=25; CHAMP1_2^+/+^ 97 ± 10 pA/pF n=15; ANOVA, mean ± SEM; Fig. S4A-B (top); Max Inward Na^+^; CHAMP1^+/+^ 81 ± 16 pA/pF n=25; CHAMP1_2^+/+^ 112 ± 15 pA/pF n=15; ANOVA, mean ± SEM; Fig. S4A-B (bottom); DIV 60-70, Max Outward K^+^; CHAMP1^+/+^ 109 ± 12 pA/pF n=18, CHAMP1_2^+/+^ 78 ± 4.3 pA/pF n=22; ANOVA, mean ± SEM; Fig. S4C-D (top); Max Inward Na^+^; CHAMP1^+/+^ 103 ± 24 pA/pF n=17, CHAMP1_2^+/+^ 89 ± 9.1 pA/pF n=22; ANOVA, mean ± SEM; Fig. S4C-D (bottom)).

Together this data suggests that CHAMP1 plays an active role in the functional development of excitatory neurons at the level of voltage-gated sodium ion channels, a key component in depolarization during neuronal firing. Other ASD and NDD related genetic mutations have been associated with alterations at the level of ion channels, including direct mutations to sodium channels^58–62^. Taken together, our results position CHAMP1 as a potential key regulator of the ion channel transitions required for proper neuronal excitability.

### CHAMP1 mutations decrease amplitude of spontaneous excitatory postsynaptic currents

Genes associated with ASD are known to regulate synaptogenesis, and altered synaptic density has been reported in ASD models^63^. Based on this, we hypothesized that CHAMP1 may contribute to the development of synaptic connections, thereby affecting cell-autonomous excitability, network connectivity, or both.

To test this, we recorded spontaneous excitatory postsynaptic currents (sEPSCs) in neurons at DIV 30-40 and 60-70 (Fig. 5; Table S5; Methods) and quantified the frequency (Hz) and amplitude (pA) of events, as well as the proportion of neurons that received at least 3 synaptic events in 2 minutes. Although the dual-SMAD inhibition protocol is designed to generate excitatory neurons, we confirmed that the synaptic events were excitatory as they were abolished in the presence of DNQX (blocks AMPA receptors) and CPP (blocks NMDA receptors) (Figure S5 A).

**Figure 5.**
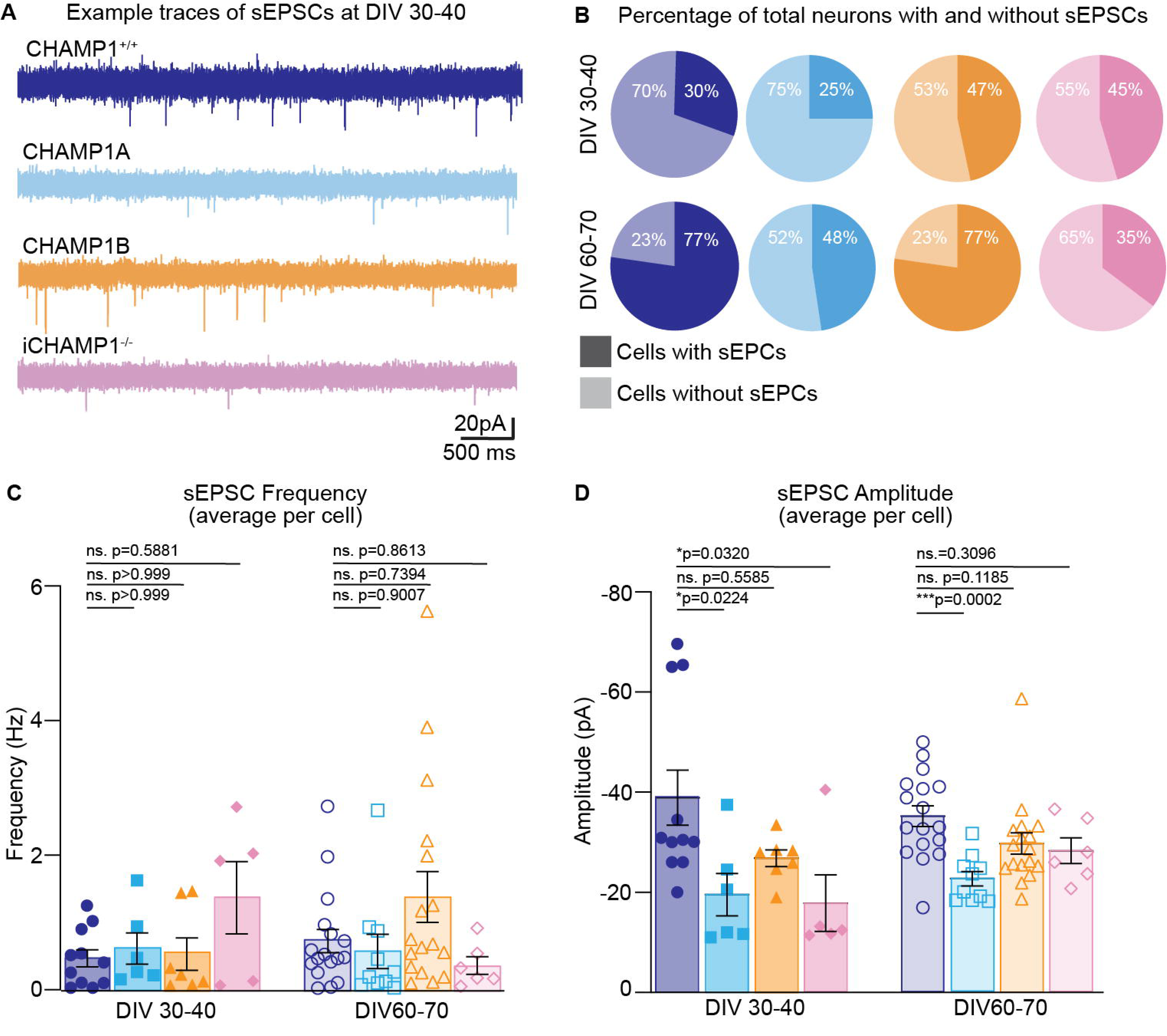
CHAMP1 mutations reduce amplitude of spontaneous excitatory postsynaptic currents. ***A***, Example traces (5 seconds) of spontaneous EPSCs in 4 cell lines at DIV 30. Neurons were held at −70mV in voltage clamp mode. ***B***, Percentage of total neurons with (darker color) and without (lighter color) sEPSCs at DIV 30-40 (top row) and DIV 60-70 (bottom row). ***C***, Average sEPSC frequency per cell in Hz at DIV 30-40 and DIV 60-70. There was no significant difference in sEPSC frequency in any group compared with CHAMP1^+/+^ within timepoints. ***D***, Average sEPSC amplitude per cell in pA at DIV 30-40 and DIV 60-70. sEPSC amplitude was significantly reduced in CHAMP1A (*p=0.0224) and iCHAMP1^-/-^ (p=0.0320) compared to CHAMP1^+/+^ at DIV 30-40. SEPSC amplitude was significantly reduced in CHAMP1A (***p=0.0002) at DIV 60-70.

We quantified the average frequency and amplitude of sEPSCs per cell and compared those data across groups. Interestingly, sEPSC frequency was comparable across genotypes at both timepoints, suggesting that presynaptic release probability was not detectably altered (DIV 30-40, CHAMP1^+/+^ 0.47 ± 0.13 Hz n=11, CHAMP1A 0.61 ± 0.23 Hz n=6, CHAMP1B 0.53 ± 0.24 Hz n=7, iCHAMP1^-/-^ 1.4 ± 0.54 Hz n=5; DIV 60-70, CHAMP1^+/+^ 0.73 ± 0.17 Hz n=17, CHAMP1A 0.57 ± 0.25 Hz n=10, CHAMP1B 1.4 ± 0.38 Hz n=17, iCHAMP1^-/-^ 0.36 ± 0.13 Hz n=6; ANOVA, mean ± SEM; Fig. 5C). In contrast, sEPSC amplitudes were significantly reduced in CHAMP1A and iCHAMP1^-/-^ neurons at DIV 30–40, with CHAMP1A deficits persisting at DIV 60–70, while CHAMP1B remained unaffected (DIV 30-40, CHAMP1^+/+^ 39 ± 5.5 pA n=11, CHAMP1A 20 ± 4.2 pA n=6, CHAMP1B 27 ± 1.7 pA n=7, iCHAMP1^-/-^ 18 ± 5.7 pA n=5; DIV 60-70, CHAMP1^+/+^ 35 ± 2.1 n=17, CHAMP1A 23 ± 1.5 n=10, CHAMP1B 30 ± 2.1 n=17, iCHAMP1^-/-^ 28 ± 2.6 pA n=6; ANOVA, mean ± SEM; Fig. 5D). The proportion of synaptically active neurons also revealed mutation-specific deficits: CHAMP1^+/+^ and CHAMP1B reached ∼77% connectivity at DIV 60–70, versus 48% (CHAMP1A) and 35% (iCHAMP1^-/-^) (Fig. 5B). Together, these data indicate CHAMP1 dosage-dependent postsynaptic dysfunction and impaired network integration during neuronal maturation. Additionally, we noted no significant difference in sEPSC frequency between CHAMP1^+/+^ and CHAMP1_2^+/+^ neurons at either timepoint (DIV 30-40, CHAMP1^+/+^ 0.47 ± 0.13 Hz n=11, CHAMP1_2^+/+^ 1.0 ± 2.6 Hz, n=11; DIV 60-70, CHAMP1^+/+^ 0.73 ± 0.17 n=17, CHAMP1_2^+/+^ 1.2 ± 0.27 n=14, ANOVA, mean ± SEM; Fig. S5).

### CHAMP1 mutations disrupt neurite outgrowth and complexity

A key aspect of neuronal maturation frequently disrupted in NDD/ASD models is dendritic arborization ^64–66^. To assess this, we immunostained iPSC-derived neurons for MAP2 at DIV 30–40 and 60–70, acquired z-stack confocal images, and generated 3D reconstructions using Imaris software (Fig. 6A-E; Methods). Sholl analysis was used to quantify branching complexity, defined as the number of intersections at 10 µm concentric rings from soma, along with total neurite length across ≥30 MAP2+ neurons per genotype/timepoint. Interestingly, at DIV 30–40, CHAMP1A neurons exhibited significantly reduced maximum Sholl intersections relative to CHAMP1^+/+^ controls, indicative of early dendritic outgrowth deficits, while CHAMP1B and iCHAMP1^-/-^ were comparable (Maximum Intersections DIV 30- 40, CHAMP1^+/+^ 5.4 ± 0.17 n=40, CHAMP1A 3.7 ± 0.28 n=34, CHAMP1B 4.7 ± 0.41 n=36, iCHAMP1^-/-^ 5.5 ± 0.21 n=40; ANOVA, mean ± SEM; Fig. 6F; Table S6). By DIV 60–70, iCHAMP1^-/-^ neurons showed the most pronounced impairment, with approximately 40% fewer maximum intersections (Maximum Intersections DIV 60-70, CHAMP1^+/+^ 6.9 ± 0.24 n=40, CHAMP1A 6.2 ± 0.58 n=44, CHAMP1B 6.3 ± 0.37 n=30, iCHAMP1^-/-^ 5.5 ± 0.28 n=40; ANOVA, mean ± SEM; Fig. 6F) and an increased average neurite length versus CHAMP1^+/+^ (Average neurite length, CHAMP1^+/+^ 26 ± 0.92 μm n=40, CHAMP1A 49 ± 4.1 μm n=44, CHAMP1B 52 ± 4.7 μm n=30, iCHAMP1^-/-^ 41 ± 3.7 μm n=40, ANOVA, mean ± SEM; Fig. 6G). CHAMP1A neurons showed a partial recovery, whereas CHAMP1B arborization remained comparable to controls. Overall, dendritic complexity increased over time in all genotypes except iCHAMP1^-/-^, indicating CHAMP1 dosage-dependent delays in dendritic maturation that parallel the electrophysiological immaturity observed in these cells. Together, these findings identify CHAMP1 as a temporal regulator of neuronal maturation, coordinating the emergence of intrinsic excitability and dendritic complexity.

**Figure 6.**
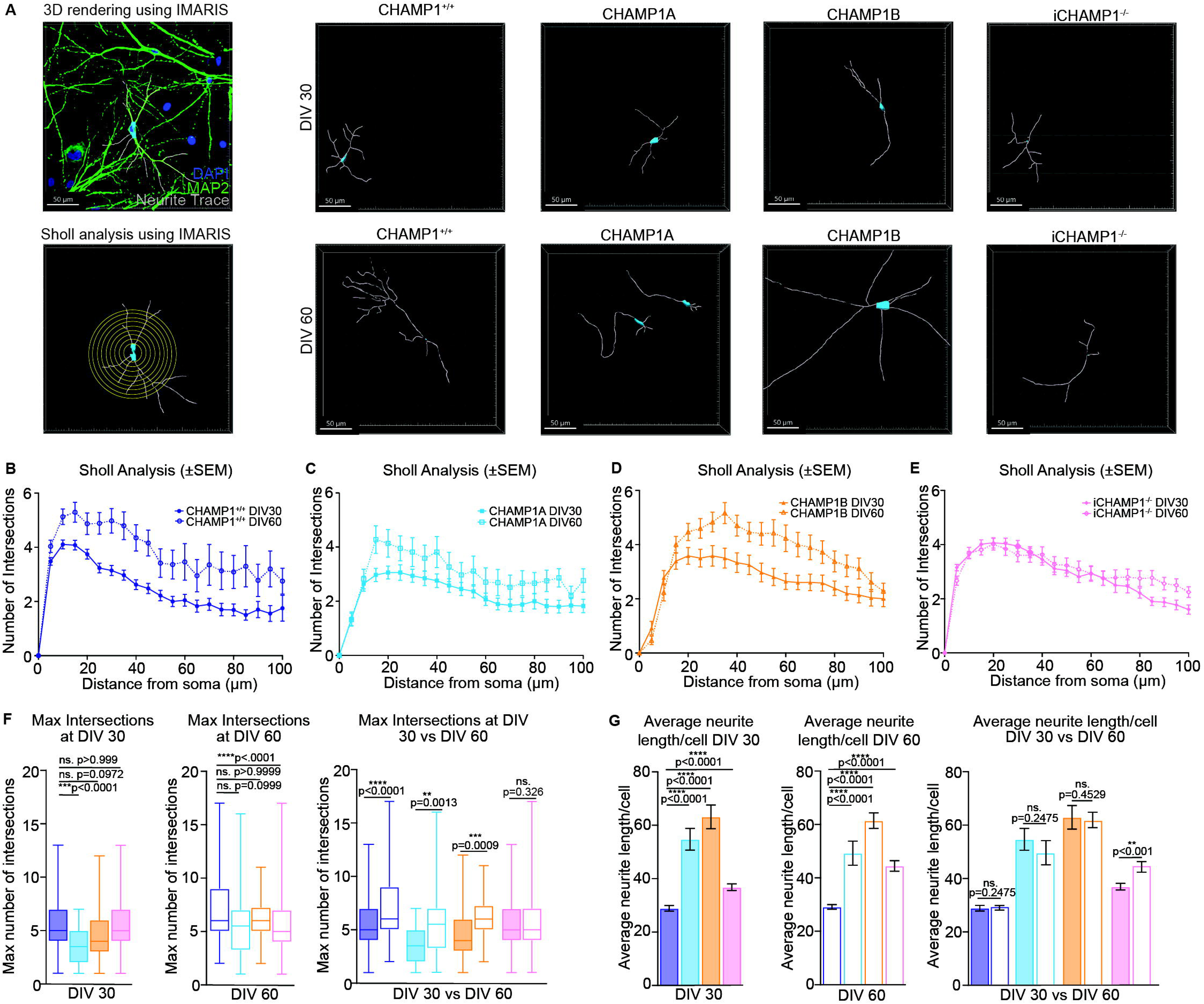
CHAMP1 mutations disrupt neurite outgrowth and branching. ***A***, 2D neurons were fixed at DIV 30 and DIV 60 and immunostained with mature neuronal marker MAP2. Z-stack images were taken on a confocal microscope for 3D reconstruction of neurites using IMARIS software. Sholl analysis quantified branch complexity by measuring the number of intersections at each concentric circle from the cell body. ***B-E***, Average number of intersections (y) plotted vs distance from soma at every 5 μm (x). Plots display within-group comparisons at DIV 30 vs DIV 60. ***F***, The maximum number of intersections was reduced in CHAMP1A neurons vs CHAMP1^+/+^ at DIV 30, and reduced in iCHAMP1^-/-^ at DIV 60. Within group comparisons reveal that CHAMP1^+/+^, CHAMP1A, and CHAMP1B increase in branch complexity between DIV 30 and DIV 60, but iCHAMP1^-/-^ does not significantly change across time. ***G***, Average neurite length (μm) was increased in all groups compared with CHAMP1^+/+^ at DIV 30 and DIV 60. This measurement did not significantly change within-group between DIV 30 and DIV 60, except for an increase in iCHAMP1^-/-^. These data suggest an aberrant growth pattern in CHAMP1 mutant neurons reflected in less complex but longer average neurites.

### CHAMP1 mutations alter gene regulatory programs during neuronal development

To determine whether these functional and structural abnormalities reflect altered developmental programs or intrinsic synaptic dysfunction, we next performed single-nucleus RNA sequencing (snRNA-seq) at DIV 30 and DIV 60 (Methods). Following quality control, a total of 140,610 high-quality nuclei were retained, with a median of 1,196 detected genes and 1,784 unique molecular identifiers (UMIs) per nucleus (Fig. S6A-B). Data from individual human iPSC lines were integrated, followed by UMAP-based clustering and cell-type annotation using established marker genes. This analysis identified four major cell types: cycling neuronal progenitors (ccNPCs), neuronal progenitor cells (NPCs), immature neurons, and mature neurons (Fig. 7A–C), supported by canonical markers (MKI67/ASPM/TOP2A in ccNPCs; NES/SOX2/VIM in NPCs; MEIS2/NCAM1/GRIA1/FOXP2 in immature neurons; and MAP2/BCL11B/SLC17A6/NRXN1 in mature neurons; Fig. 7C; Table S7).

**Figure 7.**
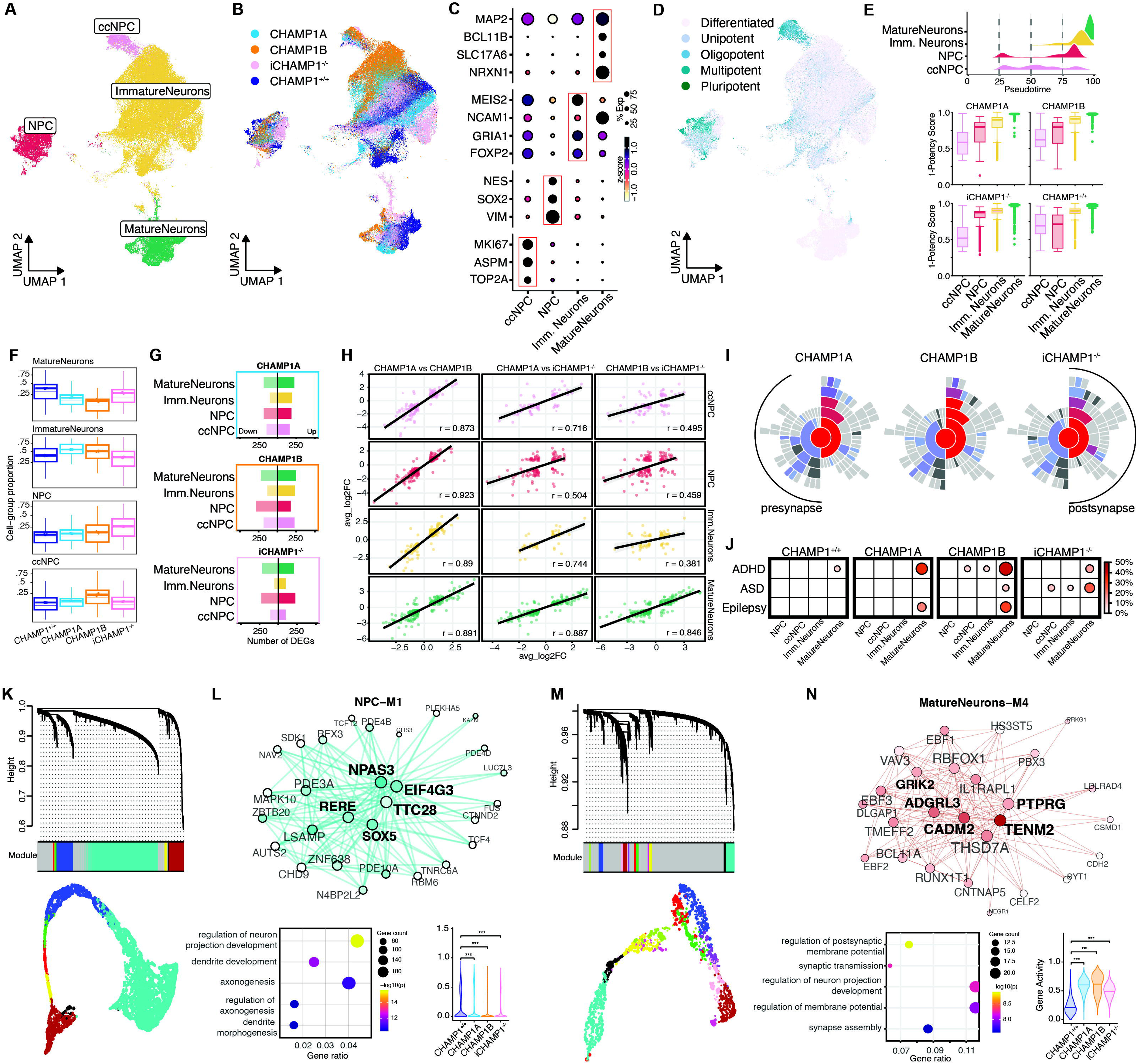
CHAMP1 mutations delay neuronal differentiation and converge on dysregulated synaptic gene networks. ***A***, UMAP projection of 140,610 integrated single nuclei from all four iPSC lines (CHAMP1^+/+^, CHAMP1A, CHAMP1B, iCHAMP1^-/-^), profiled by snRNA-seq at DIV 30 and DIV 60, colored by the four annotated cell types: cycling neural progenitor cells (ccNPCs), neural progenitor cells (NPCs), immature neurons, and mature neurons. ***B***, UMAP colored by iPSC line, showing the contribution of each genotype across cell types. ***C***, Dot plot of canonical marker genes used for cell-type annotation. Dot size indicates the percentage of nuclei expressing each gene and color indicates scaled mean expression (z-score); red boxes highlight cell-type– enriched markers. ***D***, UMAP colored by CytoTRACE2 developmental potency category, ranging from differentiated to pluripotent. ***E***, Pseudotime distribution across cell types (top) and per-cell developmental potency (1 − potency score) for each cell type, split by genotype (bottom), recovering the expected neurogenic trajectory from progenitors to mature neurons. ***F***, Proportion of each cell type per genotype, showing a reduced mature-neuron and expanded progenitor fraction in CHAMP1-mutant lines relative to CHAMP1^+/+^. ***G***, Number of differentially expressed genes (DEGs; down, left; up, right) per cell type for each mutant line relative to CHAMP1^+/+^; DEG burden is greatest in mature neurons compared with other cell-types. ***H***, Pairwise correlation of average log2 fold-changes between mutant lines within each cell type (Pearson r), showing high concordance in mature neurons (r = 0.85–0.89) and greater divergence in progenitor populations, particularly for iCHAMP1^-/-^ (r ≈ 0.46–0.50). ***I***, SynGO sunburst plots of synaptic gene-ontology enrichment among dysregulated genes for CHAMP1A, CHAMP1B, and iCHAMP1^-/-^, spanning pre- and post-synaptic compartments. ***J***, Enrichment of neurodevelopmental and psychiatric risk-gene sets (ADHD, ASD, epilepsy) among cell-types for each genotype; dot size and color indicate the percentage of cells, with the strongest enrichment in maturing and mature neurons. ***K***, Hierarchical clustering dendrogram and module assignment from co-expression network analysis (hdWGCNA) of progenitor populations (modules indicated by the color bar; trajectory shown on UMAP), together with functional enrichment of the NPC (turquoise) module and module activity across genotypes (violin; ***P < 0.01, Wilcoxon Rank Sum Test). ***L***, Co-expression network of the NPC (NPC-M1) module; node size reflects intramodular connectivity (kME) and hub genes are labeled. ***M***, hdWGCNA dendrogram and module assignment for mature neurons, with trajectory shown on UMAP. ***N***, Co-expression network of the MatureNeurons-M4 module (hub genes GRIK2, ADGRL3, CADM2, TENM2, PTPRG), its functional enrichment, and module activity across genotypes (violin; ***P < 0.01, Wilcoxon Rank Sum Test).

To investigate the differences in the neurogenic timeline, we applied trajectory inference analysis using CytoTRACE2^34^, which ordered nuclei along a continuous trajectory from high-potency cycling progenitors to low-potency mature neurons (Fig. 7D–E). Relative to CHAMP1^+/+^, CHAMP1-mutant lines showed a shifted distribution along this trajectory, with a greater proportion of nuclei retained in progenitor and immature-neuron states and a corresponding reduction in the mature-neuron compartment (Fig. 7E–F; Fig. S6C). This pattern may indicate a delayed or stalled progression toward terminal neuronal identity rather than a discrete change in cell-state composition, consistent with the delayed functional maturation observed electrophysiologically.

To quantify the transcriptional impact of each mutation, we performed differential expression analysis within each cell type (Methods). The number of differentially expressed genes (DEGs) increased with maturation and was greatest in mature neurons across all mutant lines (Fig. 7G), indicating that the transcriptional consequences of CHAMP1 loss accumulate as cells differentiate. Pairwise comparison of fold-changes for commonly differential expressed genes revealed both convergent and divergent programs: mutant lines were highly concordant in mature neurons (r = 0.85– 0.89) but diverged in progenitor populations, particularly for CHAMP1^−/−^ (Pearson’s correlation ≈ 0.46– 0.50 in ccNPCs and NPCs; Fig. 7H; Fig. S6D). Moreover, the overlap of DEGs across mutant lines further illustrates that a core set of commonly dysregulated genes is shared across all three perturbations, alongside mutation-specific components that are most pronounced in progenitor populations (Fig. S6E).

Thus, the three CHAMP1 perturbations converge on a shared neuronal endpoint while exerting mutation-specific effects at earlier developmental stages. Genes dysregulated in mutant neurons were enriched for synaptic ontologies spanning both pre- and post-synaptic compartments (Fig. 7I) and for risk genes associated with neurodevelopmental and psychiatric conditions, including ASD, ADHD, and epilepsy, with the strongest enrichment in immature and mature neurons (Fig. 7J).

To resolve the gene programs underlying these changes, we used a cell-type–resolved co-expression networks approach using hdWGCNA^45^. Within neural progenitors, we identified a module (NPC-M1) enriched for neuron projection development, dendrite development, and axonogenesis (Fig. 7K), whose hub genes included NPAS3, SOX5, RERE, AUTS2, and EIF4G3 (Fig. 7L). Interestingly, several of these hub genes are themselves high-confidence neurodevelopmental risk genes and regulators of cortical neurogenesis^67–71^. Activity of this NPC module was significantly reduced across all three CHAMP1-mutant lines relative to CHAMP1^+/+^ (Fig. 7M). Because this program is already attenuated at the progenitor stage, these data indicate that the transcriptional machinery required for neurite and axon outgrowth is downregulated early in development, providing a developmental origin for the reduced neurite branching observed morphologically and anticipating the delayed functional maturation of mutant neurons. Interestingly, a selected module in mature neurons (MatureNeurons-M4) was enriched for synaptic transmission, regulation of postsynaptic membrane potential, synapse assembly, with hub genes including GRIK2, ADGRL3, CADM2, TENM2, and PTPRG, and was likewise induced in mutant lines (Fig. 7N). These data may indicate a premature and compensatory transcriptional induction of synaptic gene programs in structurally immature neurons in which functional synaptic contacts have yet to be established. Additionally, enrichment analysis of mature neuron co-expression modules against ASD genes confirmed that MatureNeurons-M4 carries the strongest ASD signal among all mature neuron modules (Fig. S6F), further supporting its relevance to the broader neurodevelopmental risk architecture convergently affected by CHAMP1 loss.

Taken together, our single-nucleus transcriptomic analyses indicate that CHAMP1 loss perturbs the temporal program of human neurogenesis from its earliest stages, attenuating progenitor-stage pathways involved in neurite and axon outgrowth and converging on dysregulated synaptic programs in mature neurons, thereby linking proliferative and DNA repair deficits in CHAMP1-mutant progenitors to the morphological and electrophysiological phenotypes observed in differentiated neurons.

## DISCUSSION

In this study, we provide a multi-scale characterization of CHAMP1 function in human neurodevelopment, integrating cellular, electrophysiological, and single-cell transcriptomic analyses in patient-derived systems. Our findings reveal that CHAMP1 loss of function disrupts neural development across multiple layers, from impaired DNA damage responses in progenitors to altered developmental trajectories, dysregulated gene expression, and functional deficits in post-mitotic neurons, positioning CHAMP1 as a key regulator of genome stability and transcriptional control in the developing human brain.

Previous studies using NDD iPSC-derived neural progenitor cells (NPCs) have revealed abnormalities in progenitor proliferation, differentiation, and neuronal migration, supporting the notion that disruptions in early neurodevelopmental processes precede and contribute to the synaptic and connectivity defects observed across neurodevelopmental disorders^72–74^. Consistent with these observations, CHAMP1-mutant NPCs exhibited impaired proliferative capacity and defects in homologous recombination (HR)-mediated DNA repair. Specifically, CHAMP1 mutations reduced pRPA2 foci formation following CPT treatment and decreased proliferation in a genotype-dependent manner, with the most pronounced growth impairment observed in the complete knockout, suggesting increased replication stress and/or impaired cell-cycle progression. Interestingly, the HR phenotype did not follow a simple allelic dose-response relationship. CHAMP1A NPCs displayed the strongest reduction in pRPA2 foci, whereas iCHAMP1⁻^/^⁻ cells showed no significant decrease relative to CHAMP1^+/+^ controls, raising the possibility that complete CHAMP1 loss activates compensatory DNA repair mechanisms that are not engaged in heterozygous mutant cells. Recent studies in non-neuronal systems demonstrated that CHAMP1 protein-truncating variants, including Arg497*, induce HR deficiency through haploinsufficiency, with mutant proteins acting as loss-of-function alleles that do not exert dominant-negative effects on residual wild-type CHAMP1^48^. Our findings in patient-derived NPCs are broadly consistent with this model, as heterozygous CHAMP1 mutations were associated with impaired HR repair. However, the absence of a detectable HR defect in iCHAMP1⁻/⁻ NPCs suggests that haploinsufficiency alone cannot fully explain CHAMP1-dependent repair phenotypes in neural progenitors. One potential explanation is that complete CHAMP1 loss elicits cell-type-specific compensatory responses, including increased utilization of non-homologous end joining (NHEJ), alternative end joining (alt-EJ), or backup HR pathways, thereby maintaining DSB repair capacity despite the absence of CHAMP1. Such compensatory mechanisms may not be activated under heterozygous conditions or in the somatic cell systems in which CHAMP1 haploinsufficiency was originally defined. Together, these findings underscore the importance of studying CHAMP1 function in disease-relevant neural cell types and suggest that the balance between HR and alternative DNA repair pathways differs substantially between neural progenitors and non-neuronal contexts.

Previous studies using neurodevelopmental disorders iPSC-derived neuronal models have documented impairments in synaptic transmission, including defects in excitatory synaptic function and alterations in the excitatory/inhibitory balance, alongside abnormal dendritic architecture and spine morphology that contribute to aberrant neuronal connectivity^75–81^.

In our study, CHAMP1-mutant neurons exhibit depolarized resting membrane potentials, altered input resistance, reduced action potential firing, and decreased voltage-gated sodium currents at early differentiation stages, consistent with delayed acquisition of mature electrophysiological properties. These deficits partially normalize at later timepoints, supporting a model in which CHAMP1 regulates the timing rather than the absolute capacity for neuronal maturation. At the synapse, reduced sEPSC amplitude without changes in frequency suggests a postsynaptic locus of dysfunction. Morphologically, mutant neurons display reduced neurite branching but increased average neurite length, though whether this reflects a maladaptive network response to decreased early excitability or a cell-autonomous effect of CHAMP1 in post-mitotic neurons remains unclear. It is also worth noting that the reduction in sodium channel current density at DIV 60–70 in CHAMP1A and iCHAMP1^⁻/⁻^ neurons may not be fully explained by soma size, as membrane capacitance was largely comparable across genotypes at both developmental stages, with the exception of CHAMP1B neurons at DIV 60–70, which showed increased capacitance relative to CHAMP1^+/+^ controls (Table 1).

Finally, our single-nucleus transcriptomic analyses indicate that CHAMP1 mutations act primarily on the *tempo* of neuronal differentiation rather than cell fate. Across all mutant lines, nuclei occupied the same neurogenic trajectory as controls but were disproportionately retained in progenitor and immature states, with fewer cells reaching the mature-neuron compartment. Co-expression network analysis localized an early component of this defect to neural progenitors: a module enriched for neuron projection, dendrite, and axon development, anchored by hub genes NPAS3, SOX5, AUTS2, and RERE, was significantly downregulated across mutant lines before overt neuronal differentiation. This provides a developmental origin for the reduced neurite branching observed morphologically and is consistent with delayed electrophysiological maturation, positioning the progenitor transcriptome as an upstream node from which later structural and functional phenotypes emerge.

At later stages, transcriptional consequences both accumulate and converge: differentially expressed genes were most numerous in mature neurons, and while fold-change correlations among mutant lines were low in progenitors, they were markedly higher in mature neurons, indicating convergence on a shared transcriptional endpoint. These convergent programs were enriched for postsynaptic functions and for risk loci implicated in ASD, ADHD, and epilepsy, consistent with reduced sEPSC amplitude and a smaller proportion of synaptically active neurons observed electrophysiologically. Interestingly, a mature neuron module enriched for synaptic transmission and postsynaptic regulation, with hub genes GRIK2, CADM2, TENM2, and ADGRL3, was similarly upregulated across mutants. Together, these data define an early-to-late axis of neurodevelopmental dysregulation in which CHAMP1 loss first suppresses progenitor programs governing neurite and axon outgrowth, and subsequently drives induction of synaptic gene programs in neurons that have not yet acquired full structural and transcriptional maturity.

Several limitations should be acknowledged. Two-dimensional iPSC-derived monocultures lack the glial, vascular, and circuit-level context of the intact brain and cannot capture non-cell-autonomous contributions to the observed phenotypes. Additionally, the patient-derived lines differ from controls across their entire genetic backgrounds, meaning that a portion of mutation-specific variation may reflect background effects. Isogenic correction of each patient mutation, together with an expanded panel of patient lines, will be important to disentangle variant-specific from background contributions. These limitations point to clear future directions: cortical organoids and assembloids would reintroduce laminar and circuit-level context, while isogenic lines offer the most rigorous genetic control for establishing causality and defining variant-specific mechanisms.

In conclusion, our findings situate CHAMP1 within the broader genetic architecture of neurodevelopmental disorders, where diverse NDD risk genes converge on shared pathways of chromatin regulation, transcriptional control, and synaptic function^82–84^. Our data extend this principle to a gene whose canonical role lies in chromosome segregation and DNA repair: CHAMP1 loss compromises genome integrity in dividing progenitors yet ultimately converges, in post-mitotic neurons, on the synaptic and neurodevelopmental gene programs disrupted by classical NDD genes. At the same time, the mutation-specific differences we observe across repair, electrophysiological, morphological, and transcriptomic phenotypes caution against a single unifying mechanism and highlight the value of variant-resolved models for understanding, and ultimately treating, CHAMP1-associated conditions.

## Acknowledgments

Essential conceptual feedback was provided by Nitin Khandelwal, Genevieve Konopka at UCLA and Sophia Lizarraga at Brown University. Optimization of patch clamping was enabled by assistance provided by Lori McMahon at UVA. We express our gratitude to the Translational Science Lab Shared Resource, Hollings Cancer Center, Medical University of South Carolina (P30 CA138313), for their help with single cell sequencing.

## Author contributions

Conceptualization: DN, CS, MP, SB; Methodology: DN, CS, MP, LM, SB; Investigation: DN, CS, MP, SB; Formal analysis: DN, CS, EW, BG, AL; Resources: EW, BG, AL; Software: EW, BG, AL; Visualization: DN, SB, EW, BG, AL; Supervision: LM, SB; Writing: DN, CS, MP, SB

## Data availability

All processed datasets and objects generated in this study are available from the corresponding author (S.B.) upon reasonable request. Interctive single-nucleus RNA sequencing (snRNA-seq) data, including UMAP embeddings and cell-type annotations, are publicly accessible via the following link: https://biocm-nettles-et-al-neurons-cleaned.share.connect.posit.cloud/. Raw and processed sequencing data supporting the conclusions of this work have been deposited in the Gene Expression Omnibus (GEO) under accession number:

## Code availability

All custom scripts used for data analysis and figure generation are available at https://github.com/BioinformaticsMUSC/NettlesEtAl2026.

---

Table 1: Intrinsic membrane properties of CHAMP1 mutant neurons at DIV 30-40 and DIV 60-70. ANOVA compared to DIV-matched CHAMP1^+/+^. Grey shaded boxes highlight values with statistical significance, while non-shaded boxes are not significantly different compared to DIV-matched CHAMP1^+/+^.

## References

1. Hara K, Taharazako S, Ikeda M, et al. Dynamic feature of mitotic arrest deficient 2–like protein 2 (MAD2L2) and structural basis for its interaction with chromosome alignment–maintaining phosphoprotein (CAMP). Journal of Biological Chemistry. 2017;292(43):17658–17667. doi:10.1074/jbc.M117.804237

2. Itoh G, Kanno Sichiro, Uchida KSK, et al. CAMP (C13orf8, ZNF828) is a novel regulator of kinetochore-microtubule attachment: Role of CAMP in chromosome segregation. The EMBO Journal. 2011;30(1):130–144. doi:10.1038/emboj.2010.276

3. Hempel M, Cremer K, Ockeloen CW, et al. De Novo Mutations in CHAMP1 Cause Intellectual Disability with Severe Speech Impairment. The American Journal of Human Genetics. 2015;97(3):493–500. doi:10.1016/j.ajhg.2015.08.003

4. Isidor B, Küry S, Rosenfeld JA, et al. De Novo Truncating Mutations in the Kinetochore-Microtubules Attachment Gene *CHAMP1* Cause Syndromic Intellectual Disability. Human Mutation. 2016;37(4):354–358. doi:10.1002/humu.22952

5. Garrity M, Kavus H, Rojas-Vasquez M, et al. Neurodevelopmental phenotypes in individuals with pathogenic variants in CHAMP1. Cold Spring Harb Mol Case Stud. 2021;7(4):a006092. doi:10.1101/mcs.a006092

6. Levy T, Lerman B, Halpern D, et al. CHAMP1 disorder is associated with a complex neurobehavioral phenotype including autism, ADHD, repetitive behaviors and sensory symptoms. Human Molecular Genetics. 2022;31(15):2582–2594. doi:10.1093/hmg/ddac018

7. Levy T, Pichardo T, Silver H, et al. Prospective phenotyping of CHAMP1 disorder indicates that coding mutations may not act through haploinsufficiency. Hum Genet. 2023;142(9):1385–1394. doi:10.1007/s00439-023-02578-6

8. Hauf S, Watanabe Y. Kinetochore orientation in mitosis and meiosis. Cell. 2004;119(3):317–327. doi:10.1016/j.cell.2004.10.014

9. Rieder CL, Salmon ED. The vertebrate cell kinetochore and its roles during mitosis. Trends Cell Biol. 1998;8(8):310–318. doi:10.1016/s0962-8924(98)01299-9

10. Mitchison TJ. Microtubule dynamics and kinetochore function in mitosis. Annu Rev Cell Biol. 1988;4:527–549. doi:10.1146/annurev.cb.04.110188.002523

11. Mora-Bermúdez F, Kanis P, Macak D, et al. Longer metaphase and fewer chromosome segregation errors in modern human than Neanderthal brain development. Sci Adv. 2022;8(30):eabn7702. doi:10.1126/sciadv.abn7702

12. Mora-Bermúdez F, Badsha F, Kanton S, et al. Differences and similarities between human and chimpanzee neural progenitors during cerebral cortex development. Elife. 2016;5:e18683. doi:10.7554/eLife.18683

13. Degrassi F, Damizia M, Lavia P. The Mitotic Apparatus and Kinetochores in Microcephaly and Neurodevelopmental Diseases. Cells. 2019;9(1):49. doi:10.3390/cells9010049

14. Del Castillo U, Norkett R, Gelfand VI. Unconventional Roles of Cytoskeletal Mitotic Machinery in Neurodevelopment. Trends Cell Biol. 2019;29(11):901–911. doi:10.1016/j.tcb.2019.08.006

15. Vargas-Hurtado D, Brault JB, Piolot T, et al. Differences in Mitotic Spindle Architecture in Mammalian Neural Stem Cells Influence Mitotic Accuracy during Brain Development. Curr Biol. 2019;29(18):2993–3005.e9. doi:10.1016/j.cub.2019.07.061

16. Okamoto N, Tsuchiya Y, Kuki I, et al. Disturbed chromosome segregation and multipolar spindle formation in a patient with *CHAMP 1* mutation. Mol Genet Genomic Med. 2017;5(5):585–591. doi:10.1002/mgg3.303

17. Li F, Sarangi P, Iyer DR, et al. CHAMP1 binds to REV7/FANCV and promotes homologous recombination repair. Cell Reports. 2022;40(9):111297. doi:10.1016/j.celrep.2022.111297

18. Fujita H, Ikeda M, Ui A, et al. CHAMP1-POGZ counteracts the inhibitory effect of 53BP1 on homologous recombination and affects PARP inhibitor resistance. Oncogene. 2022;41(19):2706–2718. doi:10.1038/s41388-022-02299-6

19. Nagai M, Iemura K, Kikkawa T, et al. Deficiency of CHAMP1, a gene related to intellectual disability, causes impaired neuronal development and a mild behavioural phenotype. Brain Commun. 2022;4(5):fcac220. doi:10.1093/braincomms/fcac220

20. Kurganov E, Cui L, Budnik N, et al. Characterization of the functional and clinical impacts of CACNA1A missense variants found in neurodevelopmental disorders. Sci Transl Med. 2025;17(828):eadr0024. doi:10.1126/scitranslmed.adr0024

21. Chen H, LaFlamme CW, Wang YD, et al. Patient-derived models of UBA5-associated encephalopathy identify defects in neurodevelopment and highlight potential therapeutic avenues. Sci Transl Med. 2025;17(797):eadn8417. doi:10.1126/scitranslmed.adn8417

22. Rylaarsdam L, Rakotomamonjy J, Pope E, Guemez-Gamboa A. iPSC-derived models of PACS1 syndrome reveal transcriptional and functional deficits in neuron activity. Nat Commun. 2024;15(1):827. doi:10.1038/s41467-024-44989-7

23. Lizarraga SB, Ma L, Maguire AM, et al. Human neurons from Christianson syndrome iPSCs reveal mutation-specific responses to rescue strategies. Sci Transl Med. 2021;13(580):eaaw0682. doi:10.1126/scitranslmed.aaw0682

24. Niu J, Tong CK, Francois E, Frankel WN, Chung WK, Colecraft HM. Tiered modelling of a CACNA1A D1634N mutation linked to ataxia, epilepsy and cognitive deficits. Brain. Published online February 17, 2026:awag066. doi:10.1093/brain/awag066

25. Rossignoli G, Krämer K, Lugarà E, et al. Aromatic l-amino acid decarboxylase deficiency: a patient-derived neuronal model for precision therapies. Brain. 2021;144(8):2443–2456. doi:10.1093/brain/awab123

26. Dirkx N, Kaji M, De Vriendt E, et al. Kv7.2 loss-of-function causes early hyperexcitability and network remodelling. Brain. Published online June 6, 2026:awag199. doi:10.1093/brain/awag199

27. Jhanji M, York EM, Lizarraga SB. The power of human stem cell-based systems in the study of neurodevelopmental disorders. Curr Opin Neurobiol. 2024;89:102916. doi:10.1016/j.conb.2024.102916

28. Young-Pearse TL, Morrow EM. Modeling developmental neuropsychiatric disorders with iPSC technology: challenges and opportunities. Curr Opin Neurobiol. 2016;36:66–73. doi:10.1016/j.conb.2015.10.006

29. Ardhanareeswaran K, Mariani J, Coppola G, Abyzov A, Vaccarino FM. Human induced pluripotent stem cells for modelling neurodevelopmental disorders. Nat Rev Neurol. 2017;13(5):265–278. doi:10.1038/nrneurol.2017.45

30. Boulting GL, Kiskinis E, Croft GF, et al. A functionally characterized test set of human induced pluripotent stem cells. Nat Biotechnol. 2011;29(3):279–286. doi:10.1038/nbt.1783

31. Yiangou L, Grandy RA, Morell CM, et al. Method to Synchronize Cell Cycle of Human Pluripotent Stem Cells without Affecting Their Fundamental Characteristics. Stem Cell Reports. 2019;12(1):165–179. doi:10.1016/j.stemcr.2018.11.020

32. Zheng GXY, Terry JM, Belgrader P, et al. Massively parallel digital transcriptional profiling of single cells. Nat Commun. 2017;8:14049. doi:10.1038/ncomms14049

33. Germain PL, Lun A, Garcia Meixide C, Macnair W, Robinson MD. Doublet identification in single-cell sequencing data using scDblFinder. F1000Res. 2021;10:979. doi:10.12688/f1000research.73600.2

34. Kang M, Gulati GS, Brown EL, et al. Improved reconstruction of single-cell developmental potential with CytoTRACE 2. Nat Methods. 2025;22(11):2258–2263. doi:10.1038/s41592-025-02857-2

35. Mangiola S, Roth-Schulze AJ, Trussart M, et al. sccomp: Robust differential composition and variability analysis for single-cell data. Proc Natl Acad Sci U S A. 2023;120(33):e2203828120. doi:10.1073/pnas.2203828120

36. Miller SA, Policastro RA, Sriramkumar S, et al. LSD1 and Aberrant DNA Methylation Mediate Persistence of Enteroendocrine Progenitors That Support BRAF-Mutant Colorectal Cancer. Cancer Res. 2021;81(14):3791–3805. doi:10.1158/0008-5472.CAN-20-3562

37. Finak G, McDavid A, Yajima M, et al. MAST: a flexible statistical framework for assessing transcriptional changes and characterizing heterogeneity in single-cell RNA sequencing data. Genome Biol. 2015;16:278. doi:10.1186/s13059-015-0844-5

38. Granger B, Berto S. scToppR: a coding-friendly R interface to ToppGene. Bioinformatics. 2024;40(11):btae582. doi:10.1093/bioinformatics/btae582

39. Zhang MJ, Hou K, Dey KK, et al. Polygenic enrichment distinguishes disease associations of individual cells in single-cell RNA-seq data. Nat Genet. 2022;54(10):1572–1580. doi:10.1038/s41588-022-01167-z

40. Grove J, Ripke S, Als TD, et al. Identification of common genetic risk variants for autism spectrum disorder. Nat Genet. 2019;51(3):431–444. doi:10.1038/s41588-019-0344-8

41. Demontis D, Walters RK, Martin J, et al. Discovery of the first genome-wide significant risk loci for attention deficit/hyperactivity disorder. Nat Genet. 2019;51(1):63–75. doi:10.1038/s41588-018-0269-7

42. McCormack M, Gui H, Ingason A, et al. Genetic variation in CFH predicts phenytoin-induced maculopapular exanthema in European-descent patients. Neurology. 2018;90(4):e332–e341. doi:10.1212/WNL.0000000000004853

43. de Leeuw CA, Mooij JM, Heskes T, Posthuma D. MAGMA: generalized gene-set analysis of GWAS data. PLoS Comput Biol. 2015;11(4):e1004219. doi:10.1371/journal.pcbi.1004219

44. Banerjee-Basu S, Packer A. SFARI Gene: an evolving database for the autism research community. Dis Model Mech. 2010;3(3-4):133–135. doi:10.1242/dmm.005439

45. Morabito S, Reese F, Rahimzadeh N, Miyoshi E, Swarup V. hdWGCNA identifies co-expression networks in high-dimensional transcriptomics data. Cell Rep Methods. 2023;3(6):100498. doi:10.1016/j.crmeth.2023.100498

46. Andreatta M, Carmona SJ. UCell: Robust and scalable single-cell gene signature scoring. Comput Struct Biotechnol J. 2021;19:3796–3798. doi:10.1016/j.csbj.2021.06.043

47. Yu G, Wang LG, Han Y, He QY. clusterProfiler: an R package for comparing biological themes among gene clusters. OMICS. 2012;16(5):284–287. doi:10.1089/omi.2011.0118

48. Yoshizaki Y, Ouchi Y, Kurniawan D, et al. CHAMP1 premature termination codon mutations found in individuals with intellectual disability cause a homologous recombination defect through haploinsufficiency. Sci Rep. 2024;14(1):31904. doi:10.1038/s41598-024-83435-y

49. Autar K, Guo X, Rumsey JW, et al. A functional hiPSC-cortical neuron differentiation and maturation model and its application to neurological disorders. Stem Cell Reports. 2022;17(1):96–109. doi:10.1016/j.stemcr.2021.11.009

50. Tyagi S, Higerd-Rusli GP, Akin EJ, Waxman SG, Dib-Hajj SD. Sculpting excitable membranes: voltage-gated ion channel delivery and distribution. Nat Rev Neurosci. 2025;26(6):313–332. doi:10.1038/s41583-025-00917-2

51. Achkasova KA, Subbotin PV, Zhukov VV, Filat’eva AE, Tarabykin VS, Kondakova EV. Architects of the Developing Brain: Cytoskeleton-Organizing Molecules in Neurodevelopmental Disorders. Cells. 2026;15(6):537. doi:10.3390/cells15060537

52. Gunhanlar N, Shpak G, van der Kroeg M, et al. A simplified protocol for differentiation of electrophysiologically mature neuronal networks from human induced pluripotent stem cells. Mol Psychiatry. 2018;23(5):1336–1344. doi:10.1038/mp.2017.56

53. Ebert DH, Greenberg ME. Activity-dependent neuronal signalling and autism spectrum disorder. Nature. 2013;493(7432):327–337. doi:10.1038/nature11860

54. Takarae Y, Sweeney J. Neural Hyperexcitability in Autism Spectrum Disorders. Brain Sciences. 2017;7(10):129. doi:10.3390/brainsci7100129

55. Paranjapye A, Ahmad R, Su S, et al. Autism spectrum disorder risk genes have convergent effects on transcription and neuronal firing patterns in primary neurons. Genome Res. 2025;35(11):2433–2444. doi:10.1101/gr.280698.125

56. Kole MHP, Ilschner SU, Kampa BM, Williams SR, Ruben PC, Stuart GJ. Action potential generation requires a high sodium channel density in the axon initial segment. Nat Neurosci. 2008;11(2):178–186. doi:10.1038/nn2040

57. Colbert CM, Pan E. Ion channel properties underlying axonal action potential initiation in pyramidal neurons. Nat Neurosci. 2002;5(6):533–538. doi:10.1038/nn0602-857

58. Wang HG, Bavley CC, Li A, et al. Scn2a severe hypomorphic mutation decreases excitatory synaptic input and causes autism-associated behaviors. September 29, 2021. doi:10.1172/jci.insight.150698

59. Daghsni M, Rima M, Fajloun Z, et al. Autism throughout genetics: Perusal of the implication of ion channels. Brain Behav. 2018;8(8):e00978. doi:10.1002/brb3.978

60. Lee J, Ha S, Ahn J, Lee ST, Choi JR, Cheon KA. The Role of Ion Channel-Related Genes in Autism Spectrum Disorder: A Study Using Next-Generation Sequencing. Front Genet. 2021;12:595934. doi:10.3389/fgene.2021.595934

61. van Hugte EJH, Lewerissa EI, Wu KM, et al. SCN1A-deficient excitatory neuronal networks display mutation-specific phenotypes. Brain. 2023;146(12):5153–5167. doi:10.1093/brain/awad245

62. Qu G, Merchant JP, Clatot J, et al. Targeted blockade of aberrant sodium current in a stem cell-derived neuron model of SCN3A encephalopathy. Brain. 2023;147(4):1247–1263. doi:10.1093/brain/awad376

63. De Rubeis S, He X, Goldberg AP, et al. Synaptic, transcriptional and chromatin genes disrupted in autism. Nature. 2014;515(7526):209–215. doi:10.1038/nature13772

64. Durand CM, Perroy J, Loll F, et al. SHANK3 mutations identified in autism lead to modification of dendritic spine morphology via an actin-dependent mechanism. Mol Psychiatry. 2012;17(1):71–84. doi:10.1038/mp.2011.57

65. Bacova Z, Havranek T, Mihalj D, et al. Reduced Neurite Arborization in Primary Dopaminergic Neurons in Autism-Like Shank3B-Deficient Mice. Mol Neurobiol. 2025;62(5):5838–5849. doi:10.1007/s12035-024-04652-0

66. Jiang M, Ash RT, Baker SA, et al. Dendritic arborization and spine dynamics are abnormal in the mouse model of MECP2 duplication syndrome. J Neurosci. 2013;33(50):19518–19533. doi:10.1523/JNEUROSCI.1745-13.2013

67. Li Y, Fan T, Li X, et al. Npas3 deficiency impairs cortical astrogenesis and induces autistic-like behaviors. Cell Rep. 2022;40(9):111289. doi:10.1016/j.celrep.2022.111289

68. Liu JW, Li H, Zhang Y. Npas3 regulates stemness maintenance of radial glial cells and neuronal migration in the developing mouse cerebral cortex. Front Cell Neurosci. 2022;16:865681. doi:10.3389/fncel.2022.865681

69. Kwan KY, Lam MMS, Krsnik Z, Kawasawa YI, Lefebvre V, Sestan N. SOX5 postmitotically regulates migration, postmigratory differentiation, and projections of subplate and deep-layer neocortical neurons. Proc Natl Acad Sci U S A. 2008;105(41):16021–16026. doi:10.1073/pnas.0806791105

70. Fregeau B, Kim BJ, Hernández-García A, et al. De Novo Mutations of RERE Cause a Genetic Syndrome with Features that Overlap Those Associated with Proximal 1p36 Deletions. Am J Hum Genet. 2016;98(5):963–970. doi:10.1016/j.ajhg.2016.03.002

71. Oksenberg N, Ahituv N. The role of AUTS2 in neurodevelopment and human evolution. Trends Genet. 2013;29(10):600–608. doi:10.1016/j.tig.2013.08.001

72. Marchetto MC, Belinson H, Tian Y, et al. Altered proliferation and networks in neural cells derived from idiopathic autistic individuals. Mol Psychiatry. 2017;22(6):820–835. doi:10.1038/mp.2016.95

73. Gordon A, Yoon SJ, Bicks LK, et al. Developmental convergence and divergence in human stem cell models of autism. Nature. 2026;651(8106):707–719. doi:10.1038/s41586-025-10047-5

74. Ernst C. Proliferation and Differentiation Deficits are a Major Convergence Point for Neurodevelopmental Disorders. Trends Neurosci. 2016;39(5):290–299. doi:10.1016/j.tins.2016.03.001

75. Irwin SA, Patel B, Idupulapati M, et al. Abnormal dendritic spine characteristics in the temporal and visual cortices of patients with fragile-X syndrome: a quantitative examination. Am J Med Genet. 2001;98(2):161–167. doi:10.1002/1096-8628(20010115)98:2<161::aid-ajmg1025>3.0.co;2-b

76. Sichlinger L, Hausherr M, Guerrisi S, et al. Schizophrenia risk gene ZNF804A controls ribosome localization and synaptogenesis in developing human neurons. Sci Adv. 2026;12(21):eaea0755. doi:10.1126/sciadv.aea0755

77. Mariani J, Coppola G, Zhang P, et al. FOXG1-Dependent Dysregulation of GABA/Glutamate Neuron Differentiation in Autism Spectrum Disorders. Cell. 2015;162(2):375–390. doi:10.1016/j.cell.2015.06.034

78. Wen Z, Nguyen HN, Guo Z, et al. Synaptic dysregulation in a human iPS cell model of mental disorders. Nature. 2014;515(7527):414–418. doi:10.1038/nature13716

79. Shcheglovitov A, Shcheglovitova O, Yazawa M, et al. SHANK3 and IGF1 restore synaptic deficits in neurons from 22q13 deletion syndrome patients. Nature. 2013;503(7475):267–271. doi:10.1038/nature12618

80. Gouder L, Vitrac A, Goubran-Botros H, et al. Altered spinogenesis in iPSC-derived cortical neurons from patients with autism carrying de novo SHANK3 mutations. Sci Rep. 2019;9(1):94. doi:10.1038/s41598-018-36993-x

81. Kathuria A, Nowosiad P, Jagasia R, et al. Stem cell-derived neurons from autistic individuals with SHANK3 mutation show morphogenetic abnormalities during early development. Mol Psychiatry. 2018;23(3):735–746. doi:10.1038/mp.2017.185

82. Mossink B, Negwer M, Schubert D, Nadif Kasri N. The emerging role of chromatin remodelers in neurodevelopmental disorders: a developmental perspective. Cell Mol Life Sci. 2021;78(6):2517–2563. doi:10.1007/s00018-020-03714-5

83. Ronan JL, Wu W, Crabtree GR. From neural development to cognition: unexpected roles for chromatin. Nat Rev Genet. 2013;14(5):347–359. doi:10.1038/nrg3413

84. Timpano S, Picketts DJ. Neurodevelopmental Disorders Caused by Defective Chromatin Remodeling: Phenotypic Complexity Is Highlighted by a Review of ATRX Function. Front Genet. 2020;11. doi:10.3389/fgene.2020.00885

